# Nonlinear mechanics of lamin filaments and the meshwork topology build an emergent nuclear lamina

**DOI:** 10.1101/846550

**Authors:** K. Tanuj Sapra, Zhao Qin, Anna Dubrovsky-Gaupp, Ueli Aebi, Daniel J. Müller, Markus Buehler, Ohad Medalia

## Abstract

The nuclear lamina – a meshwork of intermediate filaments termed lamins – functions as a mechanotransduction interface between the extracellular matrix and the nucleus *via* the cytoskeleton. Although lamins are primarily responsible for the mechanical stability of the nucleus in multicellular organisms, *in situ* characterization of lamin filaments under tension has remained elusive. Here, we apply an integrative approach combining atomic force microscopy, cryo-electron tomography, network analysis, and molecular dynamics simulations to directly measure the mechanical response of single lamin filaments in its three-dimensional meshwork. Endogenous lamin filaments portray non-Hookean behavior – they deform reversibly under a force of a few hundred picoNewtons and stiffen at nanoNewton forces. The filaments are extensible, strong and tough, similar to natural silk and superior to the synthetic polymer Kevlar^®^. Graph theory analysis shows that the lamin meshwork is not a random arrangement of filaments but the meshwork topology follows ‘small world’ properties. Our results suggest that the lamin filaments arrange to form a robust, emergent meshwork that dictates the mechanical properties of individual lamin filaments. The combined approach provides quantitative insights into the structure-function organization of lamins *in situ*, and implies a role of meshwork topology in laminopathies.

---

During the life-cycle of a cell, the nucleus experiences large mechanical variations and has evolved to sustain large deformations and stress^1–3^. The nuclear envelope (NE), comprising the outer (ONM) and inner nuclear membrane (INM) and the lamina underlining the INM^4^, forms a rigid-elastic shell protecting the genetic material^5, 6^. The nuclear lamina – a filamentous protein meshwork at the interface of chromatin and the nuclear membrane^4, 7^ – functions as a scaffold for binding of transcription factors^8, 9^ and provides mechanical stability to the nucleus^10–13^. The principal components of the lamina in mammalian cells are mainly four lamins: lamins A and C, and lamins B1 and B2^14, 15^. Lamins are classified as type V intermediate filaments (IFs), and share a conserved tripartite domain structure with other IFs, *viz.*, a central α-helical coiled-coil rod domain flanked by a non-helical N-terminal head and an unstructured C-terminal tail domain that hosts immunoglobulin (Ig) domains^16^.

Mutations in lamins cause an important group of diseases termed laminopathies^17^ which affect the load-bearing tissues such as striated muscles leading to mechanical failure^18, 19^. It is therefore important to understand the underlying principles of lamin mechanics in health and disease^20, 21^. Intact nuclei of *X. laevis* oocytes^22^ and human fibroblasts were previously probed mechanically using the atomic force microscope (AFM)^23, 24^ to understand the role of lamina in nuclear mechanics. Micropipette aspiration has successfully provided deformation characteristics of nuclei and the NE^25, 26^. Intact cells and whole organisms have also been employed to quantify the physical properties of the nuclear lamina using stretchable substrates^10, 27^ and microfluidic devices^28^. In an attempt to measure lamin mechanics directly, a sharpened tip of an AFM cantilever was pierced inside the nucleus through the plasma and nuclear membranes^24^.

Studies performed on nucleus and entire organisms are very informative but influenced by nuclear membranes, chromatin, and surrounding cells, respectively. Direct mechanical interrogation of native lamin filaments *in situ* remains a pertinent goal towards understanding the mechanical properties of the lamina and the nucleus in health and disease^18, 29–31^. Here, using a combined *in situ* mechanical, structural and simulation approach, we characterized lamin filaments by applying point loads and measured their deformation response and apparent failure in the native meshwork (**Fig. 1**).

**Fig. 1:**
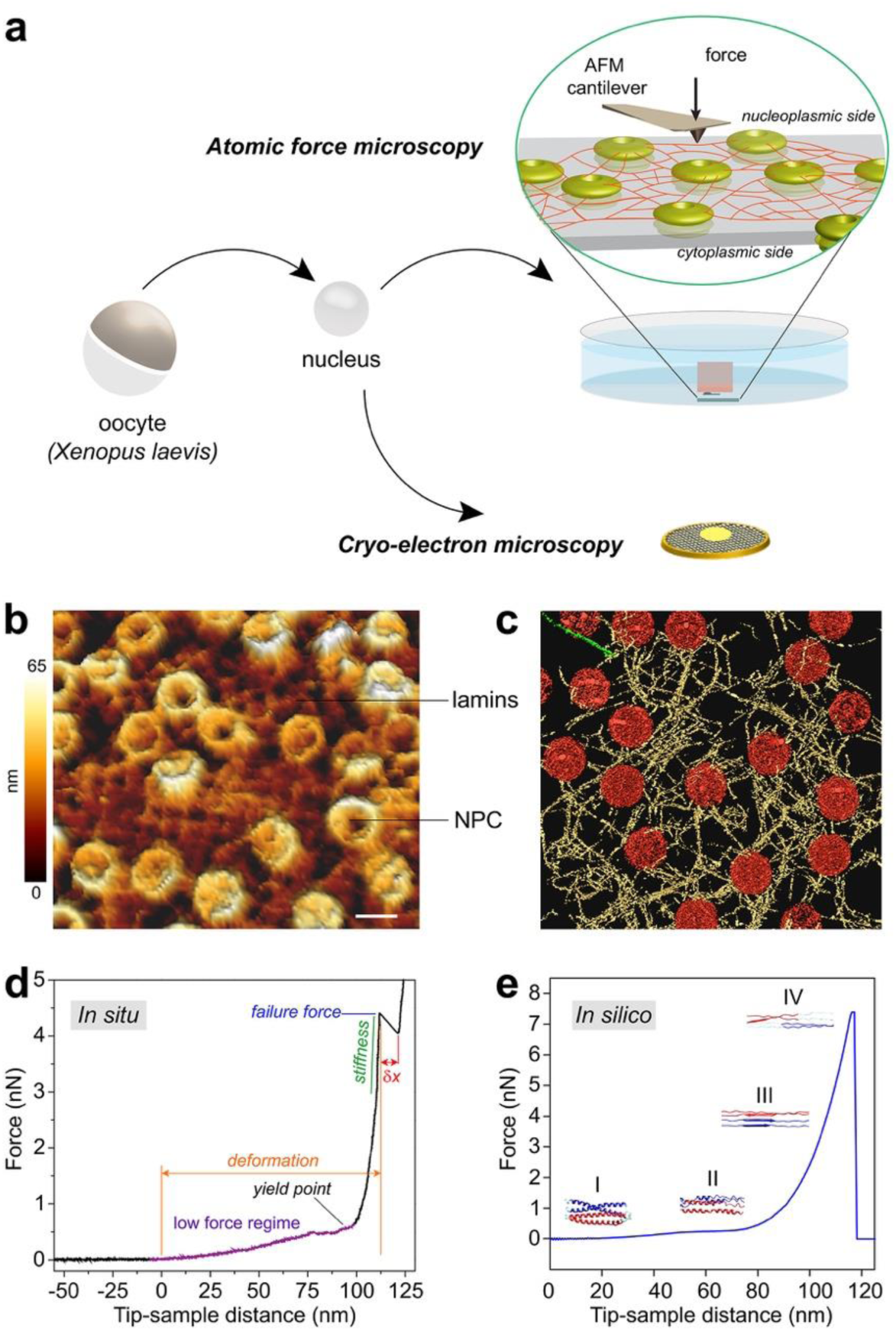
Revealing the structural mechanics of lamin filaments *in situ*. (**a**) A schematic illustration of the experimental set-up. Isolated nuclei from *X. laevis* oocytes were attached onto a poly-L-lysine-coated glass dish or carbon-coated electron microscopy grid. Next, the nuclei were manually opened by a glass capillary, and chromatin removed to expose the nucleoplasmic face. Zoom showing the cantilever tip pushing onto a lamin filament. For cryo-ET, an entire nucleus was collapsed on a grid by puncturing and removing the chromatin. (**b**) AFM imaging (nominal tip radius ≍10 nm) with a force of 0.5 nN showed areas of lamin filaments arranged orthogonally interspersed with NPCs. Scale bar, 100 nm. (**c**) Surface rendered view of a cryo-electron tomogram acquired on a spread NE of the *X. laevis* oocyte. Nuclear lamins formed a 3D meshwork of filaments (yellow) connected to NPCs (red). Field of view, 700 nm × 700 nm. (**d**) A typical FE signal showing the nonlinear behavior of a lamin filament in the meshwork. (**e**) *In silico*, a single lamin filament in the meshwork when subjected to mechanical push showed a comparable FE profile; a low force regime and a steep rise were identified indicating strain-induced stiffening leading to failure of the lamin filament. The different regions were assigned to the molecular changes in the lamin α-helical coiled coils. The yield point denotes the point of plastic or permanent deformation, *i.e.*, irreversible structural change. The similarity between the FE curves obtained *in situ* and *in silico* suggests that single lamin filaments were probed *in situ*.

Our AFM-based force-extension (FE) measurements revealed that lamin filaments deform reversibly at low loads (<500 pN), followed by stiffening that culminated in failure at >2 nN. The *in situ* mechanical behavior of lamin filaments was recapitulated *in silico* by molecular dynamics (MD) simulations of single filaments in a meshwork model^32^ derived from cryo-electron tomography (cryo-ET) of the nuclear lamina. A key finding is, that the meshwork topology influences lamin filament mechanics. The study provides a general understanding of lamin filaments and nuclear lamina mechanics relevant to laminopathies.

## Results

### *In situ* mechanics of lamin filaments

Lamin dimers assemble into staggered head-to-tail polar structures^33, 34^ that interact laterally to form tetrameric filaments^35, 36^. The filaments form a dense meshwork attached to the INM^21^ at the nuclear lamina^4, 7^. However, most lamins assemble into paracrystalline fibers in the test tube^4, 36^. Therefore, native lamin filaments can only be studied using meshwork assemblies *in situ*. First, we set out to visualize and probe lamin filaments in the nuclei of mouse embryonic fibroblasts (MEFs) and HeLa cells by AFM. Owing to the size of the mammalian nucleus (10 – 15 μm diameter), its complexity, and tight connections of the chromatin with the lamina, opening a ‘window’ into mammalian nucleus is a daunting task. For this purpose, we developed a two-step procedure: in the first step, the cell membrane was de-roofed and the nucleus exposed; in the second step, the nuclear membrane was de-roofed opening windows into the nucleus and the lamin meshwork imaged (Supplementary Figs. 1a,b, **Methods**). Nuclei from both MEFs and HeLa cells, chemically fixed (paraformaldehyde or methanol) during de-roofing, showed a filament meshwork reminiscent of the lamin meshwork observed in cryo-ET of MEF nuclei^4^ (Supplementary Figs. 1c–g). In nuclei that were not chemically fixed during de-roofing, filamentous meshwork was not observed (Supplementary Figs. 1h, i) and therefore we could not assign filaments and characterize the mechanical properties.

We therefore utilized the *Xenopus laevis* (*X. laevis*) oocyte nucleus for *in situ* AFM measurements of lamin filaments owing to its sheer size (∼400 μm diameter) and a condensed chromatin structure that is not associated with the lamina^5^. Natively assembled meshwork of single B-type lamin (lamin LIII) filaments has been visualized by electron microscopy in spread NEs of the *X. laevis* oocytes^7, 37^. In this study, the oocyte nuclei were placed on a poly-L-lysine coated glass surface to firmly attach them prior to manual opening with a sharp needle and washing the chromatin. The procedure ensured that the cytoplasmic face of the nuclear membrane (*i.e.*, the ONM) was attached to the poly-L-lysine surface and the lamin meshwork was exposed to the AFM cantilever tip. The simple isolating and spreading of NEs from the oocytes in a physiological buffer without any detergent or fixative^38^ enabled AFM and cryo-ET characterization of the lamin filaments (**Fig. 1a**). Removing chromatin by gently washing might have removed some lamin-associated proteins but retained the farnesylated lamin LIII meshwork allowing an unambiguous determination of the mechanical properties of lamins^24, 39^.

The lamina was imaged by FE-based imaging (FE-imaging) with an active closed-loop feedback, recording FE curves at 128 × 128 pixels of the sample at a force of 0.5 – 0.75 nN. As observed previously by electron microscopy methods^4, 7, 37^, lamin filaments were arranged in a meshwork exhibiting a rectangular pattern or a less organized architecture, interspersed and interacting with the NPCs (**Figs. 1b, c, Supplementary Fig. 2**). After imaging a 1 μm × 1 μm area of the lamin meshwork, the cantilever tip was positioned at random points on lamin filaments with the active closed-loop feedback, and pushed with a force of 5 – 8 nN perpendicular to the long-axis of the lamin filaments (**Figs. 1a, d**). The ONM and INM of frog oocytes are ≈50 nm apart, while NPCs are ≈90 nm tall structures^40^ with flexible lamin filaments situated on the cytoplasmic face (Supplementary Fig. 3). This enabled pushing of lamin filaments up to 100 nm towards the glass surface (**Figs. 1d, e**).

The FE curves showed an initial slow rise in the force with a plateau up to a yield point (low force regime) denoting a structural change in lamin filament where plastic or permanent deformation occurs. This was followed by considerable stiffening at larger strains indicated by a steep rise resulting in apparent failure at nanoNewton (nN) forces (high force regime) (**Fig. 1d**). FE curves obtained upon pushing the cantilever with 5 nN force on areas of the NE without filaments did not show any plateau or peaks in the low force or the high force regimes, respectively (**Supplementary Figs. 4, 5**). The nonlinear strain-stiffening behavior observed here for lamin LIII resembles previously described properties of other filamentous proteins^41–43^ including IFs^44, 45^, and rather reflects a general mechanism of material deformation and failure^46^. Strain hardening has also been reported for reconstituted lamin B networks and is dramatically higher than observed in networks of keratin, vimentin or F-actin^47^.

To elucidate the structural intermediates during the mechanical pushing of a lamin filament, we performed molecular dynamic (MD) simulations on single filaments in a meshwork model based on cryo-electron tomograms of the nuclear laminae of *X. laevis* oocytes and MEFs (see **Methods**). Mechanical behavior of single lamin filaments was simulated by applying an out-of-plane pushing force on single points in the meshwork model. Similarities between the AFM and the *in silico* FE curves suggest that we probed single lamin filaments *in situ* (**Figs. 1d, e**). The simulations showed that the plateau in the low force regime in the FE curves is due to partial unfolding of the α-helical coiled-coil domains. However, sliding between lamin dimers cannot be excluded although direct interactions of the farnesylated lamin tails with the INM makes this less likely. It should be noted that in MD simulations, which are performed at a few orders of magnitude higher speeds than experiments, we did not observe clear sliding. Sliding may occur more frequently at lower speeds (as in experiments) as at higher speeds (>>1 μm s^−1^) sliding requires much larger forces than unfolding coiled-coil α-helices thereby making unfolding the dominant event^48^. The inter-connectedness of the filaments at junctions may also resist sliding and promote strain-induced stiffening. The stiffening at high force represents the transition of α-helical coiled-coils to β-sheets followed by failure. The force tolerated by a filament before failure *in silico* was comparable to that measured *in situ*. Simulation analysis indicated the FE curve profiles did not depend on the size of the probe used to push the lamin filaments and the transition from α-helix to β-sheet occurred through similar structural intermediates (Supplementary Fig. 6a-c). It is suggested that the mechanical reaction of lamin filaments is a robust characteristic and may be key to its function during cell migration through narrow crevices^28^ or during nuclear envelope breakdown^13, 47^ when the nucleus experiences loads.

Single intermediate filaments when pushed perpendicular to the long axis^49^, stretched along the long axis^50, 51^ or dragged on a surface^52^ were able to withstand nN forces and the FE curves showed low and high force regimes. Previous MD simulations of stretching lamins in an orthogonal meshwork^32^ (Supplementary Fig. 7) also showed nonlinear stress-strain profiles and failure forces similar to those observed by mechanical pushing of lamin filaments *in situ* (**Fig. 1d**).

### Detailed mechanical behaviour of lamin filaments

The FE curves recorded *in situ* showed a plateau (15-35%) in the low force regime (**Fig. 2a, Supplementary Table 1**) at a force *F*_low_ ≈ 0.3 nN (**Fig. 2b**). The plateau presumably denotes unfolding and sliding of an α-helical coiled-coil, preceding its conversion into stiffer β-sheets^48^ (**Fig. 1e**, Supplementary Fig. 6). The ‘soft’ coiled-coils (*κ*_low_ ≈ 0.008 nN nm^−1^, (**Fig. 2c**)) experienced entropic stretching up to a deformation *d*_low_ ≈ 54 nm (**Fig. 2d**). An engineering strain of ≈126% (**Supplementary Note 1**) before stiffening is observed, similar to other semi-flexible filaments whose persistence and contour lengths are of similar magnitude^4, 53^. Unfolding of α-helical coiled-coils before transition into β-sheet is a well-known phenomenon^54^, and has been suggested for vimentin^55^, fibrin and fibrinogen during blood clotting^43, 56^.

**Fig. 2:**
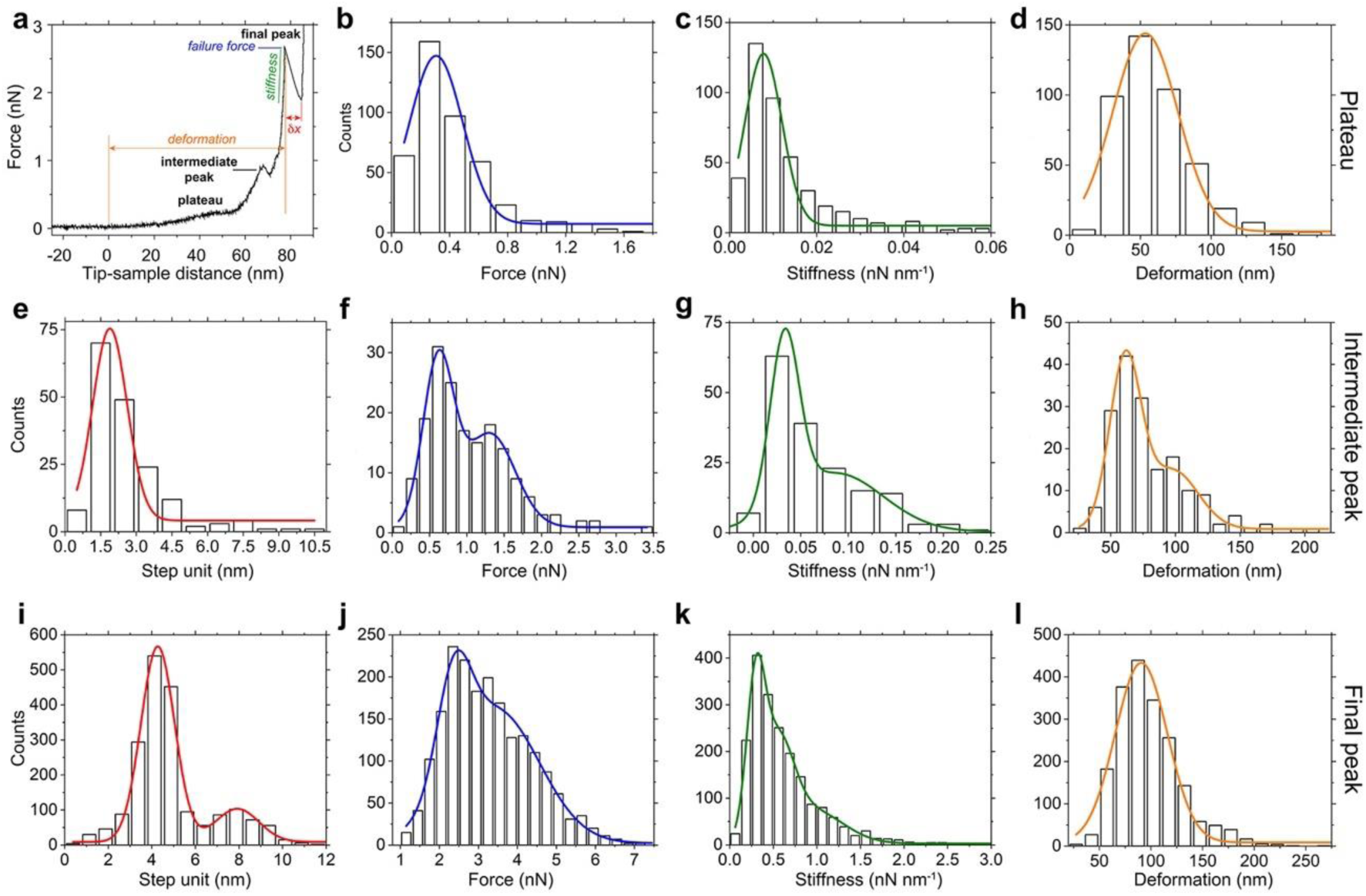
Mechanical characteristics of lamin filaments. (**a**) An FE curve showing a characteristic plateau and an intermediate peak preceding the final peak at high force (≍3 nN). (**b**) The plateau occurred at a force of 0.30 ± 0.20 nN (average ± standard deviation calculated from FWHM). (**c**) The region showed a stiffness of 0.0078 ± 0.0043 nN nm^−1^ (*n* = 432), and (**d**) a deformation of 54 ± 23 nm (*n* = 432). (**e**) The step unit of the intermediate peak was 1.9 ± 0.7 nm at a force of 0.60 ± 0.20 nN (*n* = 175) (**f**). (**g**) A stiffness of 0.033 ± 0.015 nN nm^−1^ (*n* = 172) was 5-fold more than that of the structural intermediate at the plateau. (**h**) A deformation of 61 ± 12 nm (*n* = 173) was marginally larger than that at the plateau. The shoulders in the histograms, (**f**) force (1.3 ± 0.3 nN), (**g**) stiffness (0.086 ± 0.050 nN nm^−1^), and (**h**) deformation (98 ± 20 nm) indicate a continuum of the stiffening process. (**i**) Step unit histograms (4.3 ± 0.80 nm; 7.9 ± 1.0 nm; *n* = 1946); (**j**) failure force (2.4 ± 0.40 nN; 3.5 ± 1.1 nN; *n* = 1946); (**k**) stiffness (0.30 ± 0.11 nN nm^−1^; 0.54 ± 0.19 nN nm^−1^; 0.95 ± 0.34 nN nm^−1^; *n* = 1946), and (**l**) deformation (91 ± 26 nm; *n* = 1946) of the final peak data pooled from all speeds (Supplementary Fig. 9).

Beyond the yield point and prior to the detected failure, an intermediate peak (4-16%) of step unit ≈1.9 nm was observed between 1.0 – 1.5 nN (**Fig. 2e-h, Supplementary Table 1**). A step unit of ≈1.9 nm is similar to the diameter of individual lamin dimers (≈1.7 nm)^57^ (Supplementary Fig. 8). Thus, we deduced that the intermediate peak presumbaly denotes the rupture of dimer-dimer interactions during the transformation from folded α-helices to β-sheets.

Next, in the high force regime beyond the yield point (**Fig. 1d**), lamin filaments showed plastic deformation. Plastic deformation was not observed upon indentating nuclei at increasing loading rates most likely because of limited indentation depths hindering direct probing of the lamin meshwork and interference from the NE^25^. The FE curves showed an abrupt drop in force with step units ∂*x*_high_ ≈ 4 or 8 nm (**Fig. 2i**), between *F*_high_ ≈ 1.5 – 5.0 nN (**Fig. 2j**). Cryo-ET analysis of the oocyte NE confirmed that lamin tetramers of diameter 3.8 nm associate laterally to form filaments of diameter 7 nm (Supplementary Fig. 8a arrowheads, 8c)^4, 7^.

MD simulations suggest that *in situ* filament stiffening, *κ*_high_ > 0.3 nN nm^−1^ at high forces (**Fig. 2k**), is caused by α-helix coiled-coil unfolding and transitioning to β-sheets. With a capacity to withstand an average deformation *d*_high_ ≈ 91 nm (up to 200 nm) (**Fig. 2l**), lamin filaments showed a remarkable engineering strain of *ε* ≈ 250% (**Supplementary Note 1**) similar to the average strain of ∼250% observed for single IFs^52, 58^. The strain of lamin filaments measured here is higher than that of collagen (12%), α-keratin (45%), elastin (150%)^59^, but comparable to desmin (240%)^49^, vimentin (205%)^51^ and hagfish slime (220%)^45^.

Increasing the loading rate over two orders of magnitude did not vary the step unit, force or deformation of lamin filaments (Supplementary Fig. 9). Although a hysteresis was observed between the pushing and relaxing FE curves suggesting a viscous contribution (**Fig. 3a**), lamin filaments stiffened to a similar extent at increasing loading rates. This indicates that no viscous structural element contributed to filament mechanics (Supplementary Fig. 10) suggesting that the energy is stored in the β-sheet structure^55^. Interestingly, we observed a direct correlation between the force a filament was able to tolerate before its apparent failure and the stiffness, implying that as a filament stiffened, its capacity to bear load also increased (Supplementary Fig. 11).

**Fig. 3:**
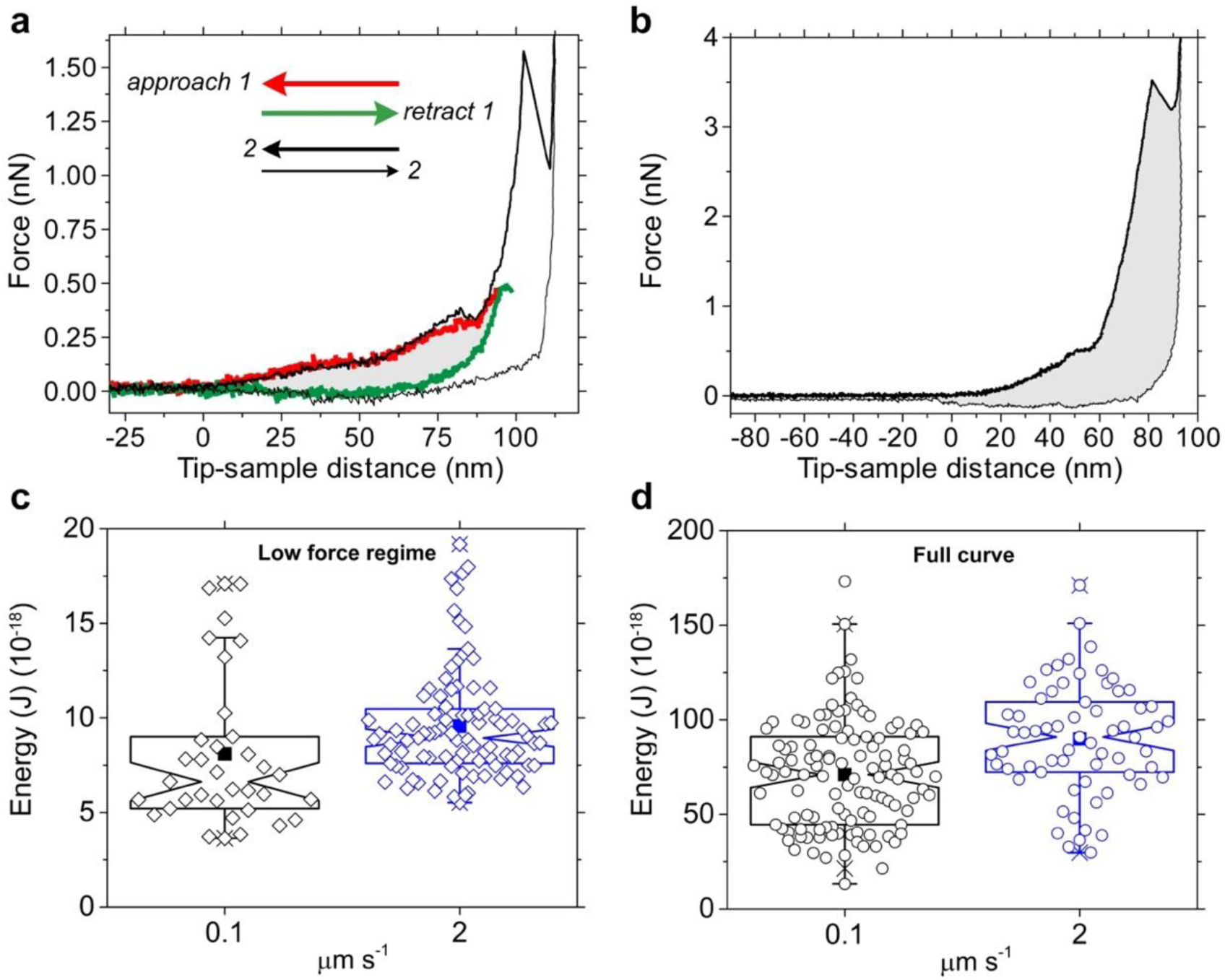
Lamin filaments absorb energy under continuous applied force. (**a**) To estimate the energy absorbed during the initial stretching of a lamin (low force regime), a repetitive force protocol was used. A force of 0.5 nN was applied (approach, red curve) and released (retract, green curve); the area between the red and green curves (grey) gives the energy absorbed by the lamin structure. The same filament was subjected to a higher force until failure was detected (bold black curve). The plateau in the low force regime was reversible (compare black and red curves) in majority (89%) of the FE curves (*n* = 133). The reversibility shows that the cantilever tip could be stably and reproducibly placed on a lamin filament, and that a single filament was interrogated. (**b**) The area (shaded grey) between the approach curve (bold black) and the retract curve (thin black) denotes the energy dissipated during the entire process of failure. (**c, d**) The energy absorbed in the low force regime was ≈10^−17^ J (*n* = 40 at 0.1 µm s^−1^, *n* = 93 at 2 µm s^−1^), an order of magnitude lower than that absorbed during the irreversible failure ≈10^−16^ J (*n* = 115 at 0.1 µm s^−1^, *n* = 65 at 2 µm s^−1^).

Following the first peak, ≈60% of the FE curves also showed a second peak with a step unit of ≈4 nm (Supplementary Figs. 9, 12), while additional peaks were seldom seen as well (Supplementary Fig. 13). Akin to the first peak, the second peak presumably indicates the failure of an additional lamin tetramer that was also detected by cryo-ET (Supplementary Fig. 8c). Alternatively, these peaks may detect interactions between lamins and other binding proteins. Control experiments however showed that the forces measured were specific to lamins since pushing directly on nuclear membranes produced FE curves without any prominent peaks (Supplementary Fig. 4). Moreover, characteristic force response of lamin LIII filaments was still obtained after nuclease treatment, and the measured parameter values did not change by incubating the NEs with the nuclease (Supplementary Fig. 14). Since nanoNewton forces are not typical of protein-protein interactions, and filaments were pushed at many hundreds of positions, we reckon that the measured parameters are predominantly of the lamin LIII filaments.

**Fig. 4:**
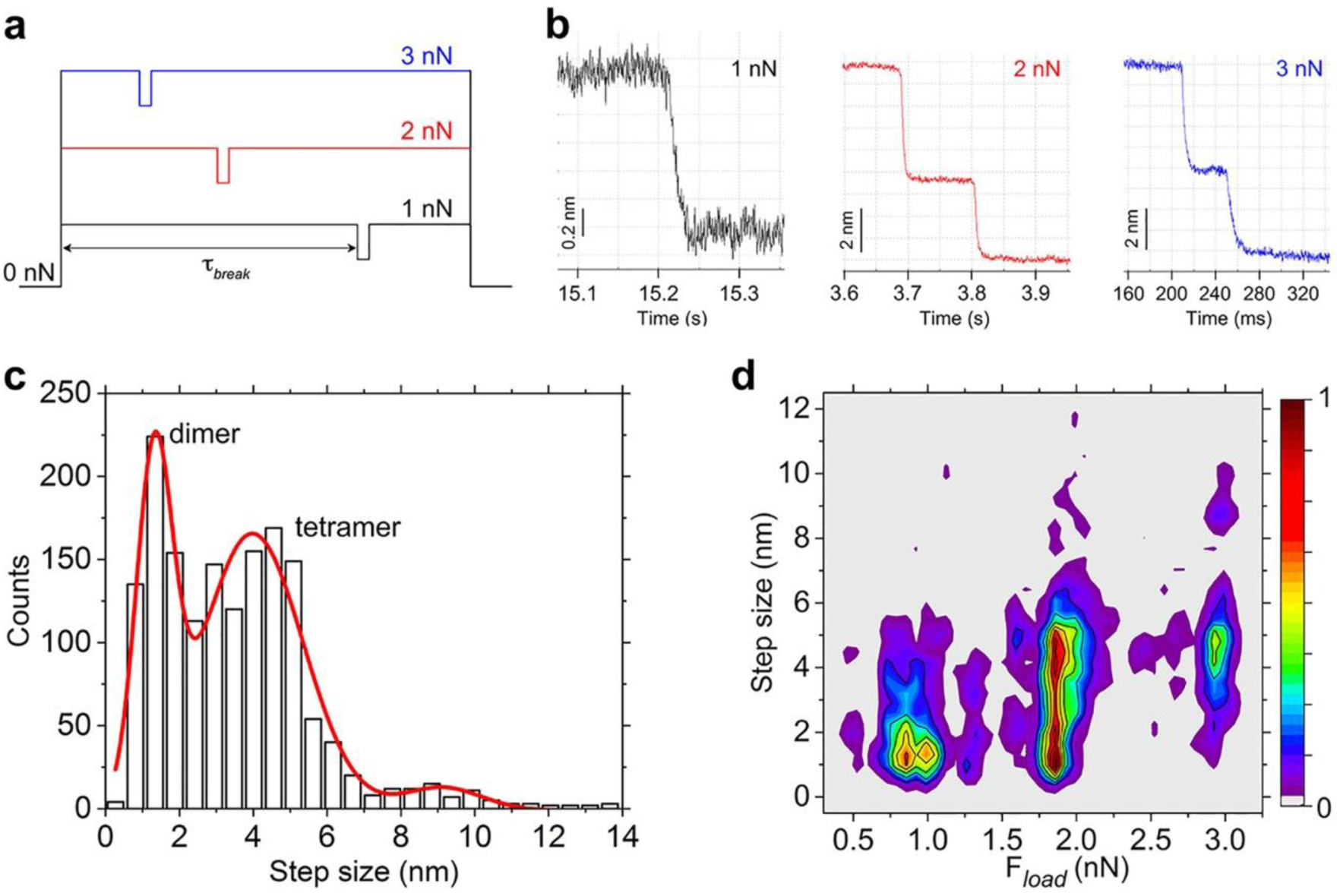
Discrete failure-steps of lamin filaments at constant loads (force clamp). (**a**) Lamin filaments were subjected to constant loads (*F*_load_) ranging from 0.75 nN to 3 nN. (**b**) Typical signals denoting the molecular alterations in lamin filaments under constant loads (*F*_load_) of 1 nN, 2 nN and 3 nN. (**c**) Failure of lamin filaments at constant *F*_load_ occurred in discrete steps of 1.3 ± 0.5 nm, 4.0 ± 1.4 nm or rarely 9.0 ± 1.1 nm (*n* = 1569). (**d**) A density map showing that the main population consists of steps ≈1 nm at low loads (≤1 nN) increasing to ≈4 nm at high loads (≥3 nN) with a transition observed at 2 nN. The 1.3 nm step is attributed to the mechanical rupture of the α-helix coiled-coil (diameter ≈ 1.2 – 1.7 nm)^4, 57^. The 4 nm steps denote the failure of stiffened tetramers at a high force. The color scale denotes normalized densities of the populations.

### Lamins are tough and act as shock absorbers

The lamin meshwork is suggested to function as a shock absorber protecting the nuclear contents from external mechanical forces^5^. A question we asked was: is shock absorption an emergent property of the meshwork or, is a single filament capable of absorbing shocks too? We directly measured the energy absorbing capacity of lamin filaments assembled *in situ*. For this, we applied a repetitive force protocol to measure the energy dissipated in the low force regime (plateau) before stiffening. The first pushing step (0.5 nN) of the protocol showed a plateau; however, the relaxation curve did not show a plateau (described previously for perfect spring proteins like myosin^60^). The second pushing step showed the plateau again resulting in a hysteresis between the pushing and the relaxation curves (**Fig. 3a**). The recurrence of the plateau suggests that either the refolding or the spring-like sliding of the α-helical coiled-coil^61^ is reversible and occurs within ≈500 ms (see **Methods**). Interestingly, the plateau force did not depend on the loading rate (Supplementary Fig. 15) and did not recur after filaments had experienced nanoNewton forces up to the apparent failure.

**Fig. 5:**
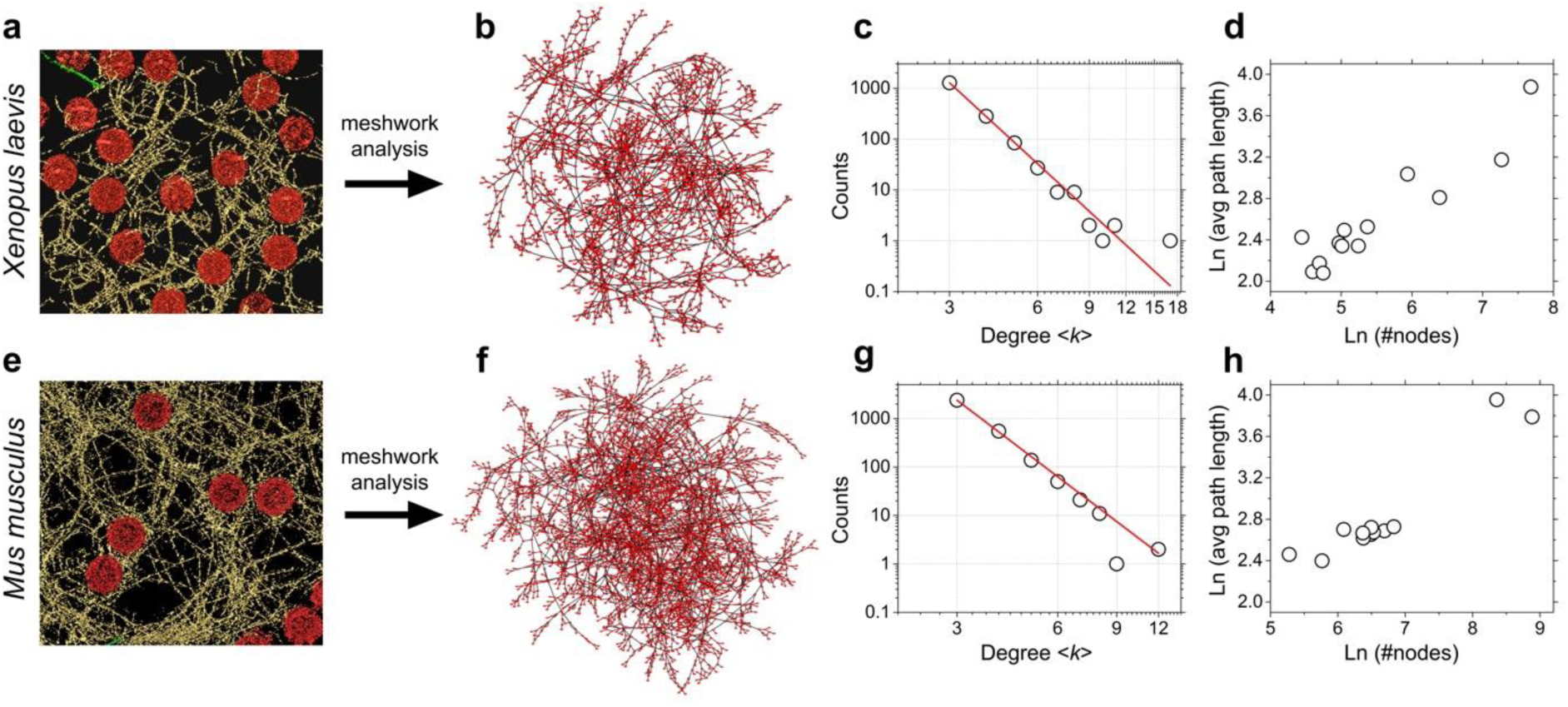
Nuclear lamin meshwork of *X. leavis* and MEF show similar topology. The 3D lamin meshworks as viewed by cryo-ET of (**a**) *X. laevis* oocyte NE (Fig. 1b) and (**e**) MEF NE (adapted from ref. *4*; field of view, 1000 nm × 1000 nm) were analyzed to create undirected graphs (**b**, **f**). Nuclear lamins formed a 3D meshwork of filaments (yellow) connected to NPCs (red). The red dots in the graphs denote the nodes or the vertices (coordinates of filament interactions or crossovers), defined as the points where adjacent lamin filaments (grey connecting lines) appeared closer than 1.3 nm to each other. (**c**, **g**) Degree distribution of both the lamin meshworks revealed a Power-law behavior with an index λ ≈ 5.6, and (**d**, **h**) exhibited ‘small world’ characteristics where the average path-lengths scale with the meshwork size.

The energy attributed to the hysteresis in the low force regime was determined to be ≈10^−17^ J (**Fig. 3a, c**) increasing to ≈10^−16^ J (10^5^ *k*_B_T) up to the failure (**Fig. 3b, d**). The energies did not change with loading rates providing evidence that lamin filaments in healthy state are ductile and not brittle^62^. A back-of-the-envelope calculation suggests that the total energy absorbed by the filaments during pushing is equivalent to that required for breaking ≈170 C-C bonds (1 C-C ≈ 5.8 × 10^−19^ J). However, the measured failure force (*F*_high_) between 2 nN – 5 nN is similar to the force required to break a single C-C bond^63^. Increasing the pushing speed of the cantilever 20-fold, and hence the kinetic energy imparted by the cantilever 400-fold, did not change the energy dissipated by lamin filaments. We therefore suggest, that the energy provided by the cantilever is not utilized for breaking covalent bonds but is expended in disrupting non-covalent interactions (sum of charges, van der Waals forces, and hydrogen bond interactions) involved in the unfolding of protein structures, including the force-induced α-helix to β-sheet transformation and the stick-slipping process of the β-sheets under tension^48^.

Based on the total energy, the tensile toughness, *T*, of a lamin filament (*r* ≈ 2 nm) is estimated to be ≈147 MJ m^−3^ (volume, *V* ≈ 679 nm^3^) or ≈10^5^ J kg^−1^. Remarkably, the toughness of a lamin filament is superior to that of high-tensile steel (6 MJ m^−3^), carbon fibre (25 MJ m^−3^) and Kevlar 49 fibre (50 MJ m^−3^), much higher than the toughness of natural materials such as elastin (2 MJ m^−3^), resilin (4 MJ m^−3^), tendon collagen (7.5 MJ m^−3^), and at par with that of wool (60 MJ m^−3^), *Bombyx mori* cocoon silk (70 MJ m^−3^), nylon (80 MJ m^−3^), *A. diadematus* dragline (160 MJ m^−3^) and viscid (150 MJ m^−3^) silks^64^. The strong, tough and extensible nature of lamin filaments offer promising possibilities for engineering lamin-based materials.

### Lamin filaments can withstand constant forces upto nanoNewtons

Is the mechanical behavior of lamin filaments peculiar to force loading at a constant velocity or could it be recaptured at constant loads too? To answer this, we subjected lamin filaments in the meshwork to constant loads (*F*_load_) of 0.75 – 3.0 nN. As in constant velocity experiments, discrete steps of 1.3 nm (*F*_load_ ≤ 1 nN), 4 nm or 8 nm (*F*_load_ ≥ 2 nN) were detected (**Fig. 4**).

The lifetime, *τ* _break_, of a few hundred milliseconds measured for the α-helical coiled-coils at *F*_load_ ≤ 1 nN suggested that it is the first buffer against mechanical shocks and acts as a sacrificial unit at lower forces. The coiled-coil structure absorbs the kinetic energy thereby preventing the force propagation to further regions of the network. As the force is increased to 3 nN, the α-helix to β-sheet transition increases the load bearing capacity of the filament because of stiffening, and the failure requires tens of milliseconds (Supplementary Fig. 16). Local stiffening and failure at nanoNewton forces may serve as an efficient mechanism for preventing breakage of other filaments in the meshwork thereby preventing a catastrophic meshwork failure^32^.

**Fig. 6.**
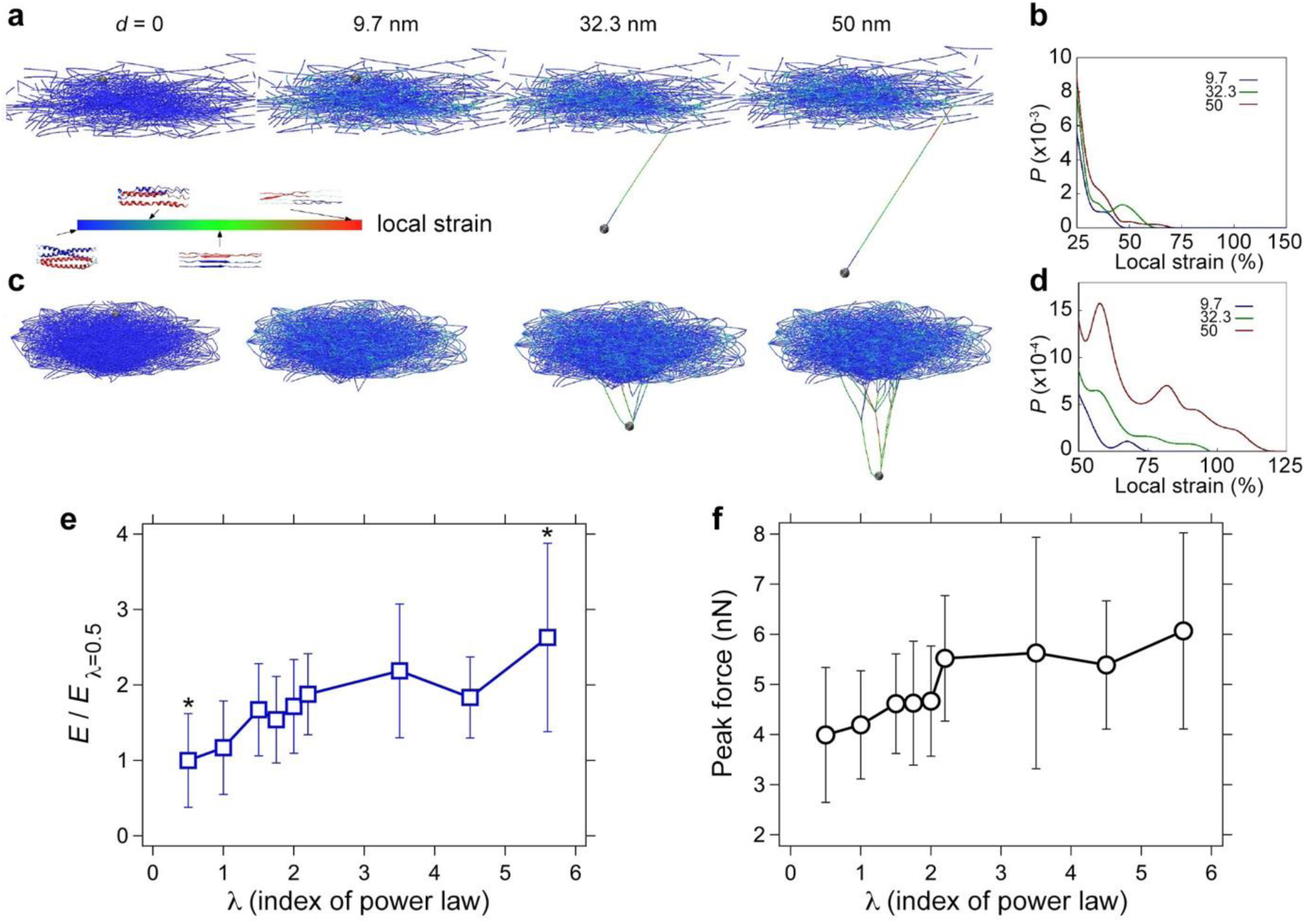
Meshwork topology (λ) influences toughness and strength of lamin filaments. Snapshots from simulations of mechanical pushing of single lamin filaments in meshworks of two different node connectivities: (**a**) λ = 0.5 and, (**c**) λ = 5.6. The meshwork of λ = 0.5 was composed of many single filaments connected to a few heavy nodes, while the meshwork of λ = 5.6 was composed of well-connected filaments with a balance of light nodes, intermediate nodes and a few heavy nodes. The color scale denotes the local strain at different deformations, *d*, of the filament. (**b, d**) Probability distributions of the local strain within each filament segment measured at increasing deformations (corresponding to snapshots in panels **a**, **c**) for the two meshworks. At λ = 0.5, the filament being pushed ruptured from one end at an early stage (*d* < 32.3 nm), the other end was stretched and unraveled until the heavy node and the strain in the filament increased to cause rupture. At λ = 5.6, increasing deformation caused a large strain re-distribution. The meshwork with λ = 5.6 was associated with a much larger cohesive zone that enabled to dissipate more deformation energy before rupture (see **Methods**). Lamin filaments in meshworks with higher λ values showed increased (**e**) toughness (*E*) and (**f**) mean strengths as compared to filaments in meshworks with smaller λ. * in (**e**) signifies *p* < 0.05.

### Meshwork topology influences lamin mechanics

Visually, the lamin meshwork appears as a random arrangement of filaments. To test if this is indeed the case, we set: (i) to compare the design features of the lamin meshwork in different species; (ii) to decipher the influence of meshwork topology on lamin filament mechanics. To address these aims, in a first step we employed cryo-ET for visualizing the nuclear laminae of *X. laevis* oocyte (**Fig. 5a**) and mouse embryonic fibroblasts (MEFs) (**Fig. 5e**, cryo-ET data from ref. ^24^). Next, we applied graph theory^65^ to compare and quantitate the meshwork topologies. To this end, the meshworks were converted to undirected 3D graphs; lamin filaments formed the links between the vertices representing the physical connections between filaments (**Figs. 5b, f**). Interestingly, the lamin meshworks of both, *X. laevis* oocyte and mammalian nuclei, exhibit similar topological features. The average degree of connectivity, <*k*>, of both the meshworks is 3.3. The degree (*k*) distribution of both the meshworks follows a Power-law [*P*(*k*) ∼ *k* ^−λ^] with a scaling factor λ > 5. In other words, the meshworks consist of many nodes with connectivity 3 or 4 and a minor population of hubs of high degrees (5 – 17) (**Fig. 5c, g**). It is noteworthy that despite the difference in lamin-types (B-type lamin LIII in *X. laevis* oocytes, and A-type and B-type MEF lamins), the lamin meshworks of both species share common topological features. Furthermore, in both meshworks the average path-length scales with the meshwork size, *i.e.*, distance scales with the number of nodes (**Fig. 5d, h**). This ‘small-world’ property of the lamin meshwork, defined simply as the nodes connected through the shortest distances, is similar to that of a power grid network^66^ and points to the importance of hubs to the integrity of the meshwork. Although we cannot elucidate the structural identity of the lamin connections, cross-linking mass spectroscopy studies suggest that electrostatic interactions between the unstructured head and tail domains of adjacent lamin dimers may drive meshwork assembly^61^.

Mutations in lamins and NE proteins^67^ are suggested to influence the mechanical properties of the nucleus^68, 69^. We conjectured that the meshwork topology may influence the mechanical properties of the lamina. To test this hypothesis, we performed MD simulations on single filaments in meshwork models of different topologies (Supplementary Fig. 17). Meshwork topologies were created by changing the λ values between 0.5 – 5.6; filaments in a meshwork of λ = 0.5 were the least connected and λ = 5.6 were the most connected (see **Methods**). For all meshwork topologies, the simulated profiles of the FE curves resembled those obtained *in situ* (**Fig. 1c**).

**Fig. 7:**
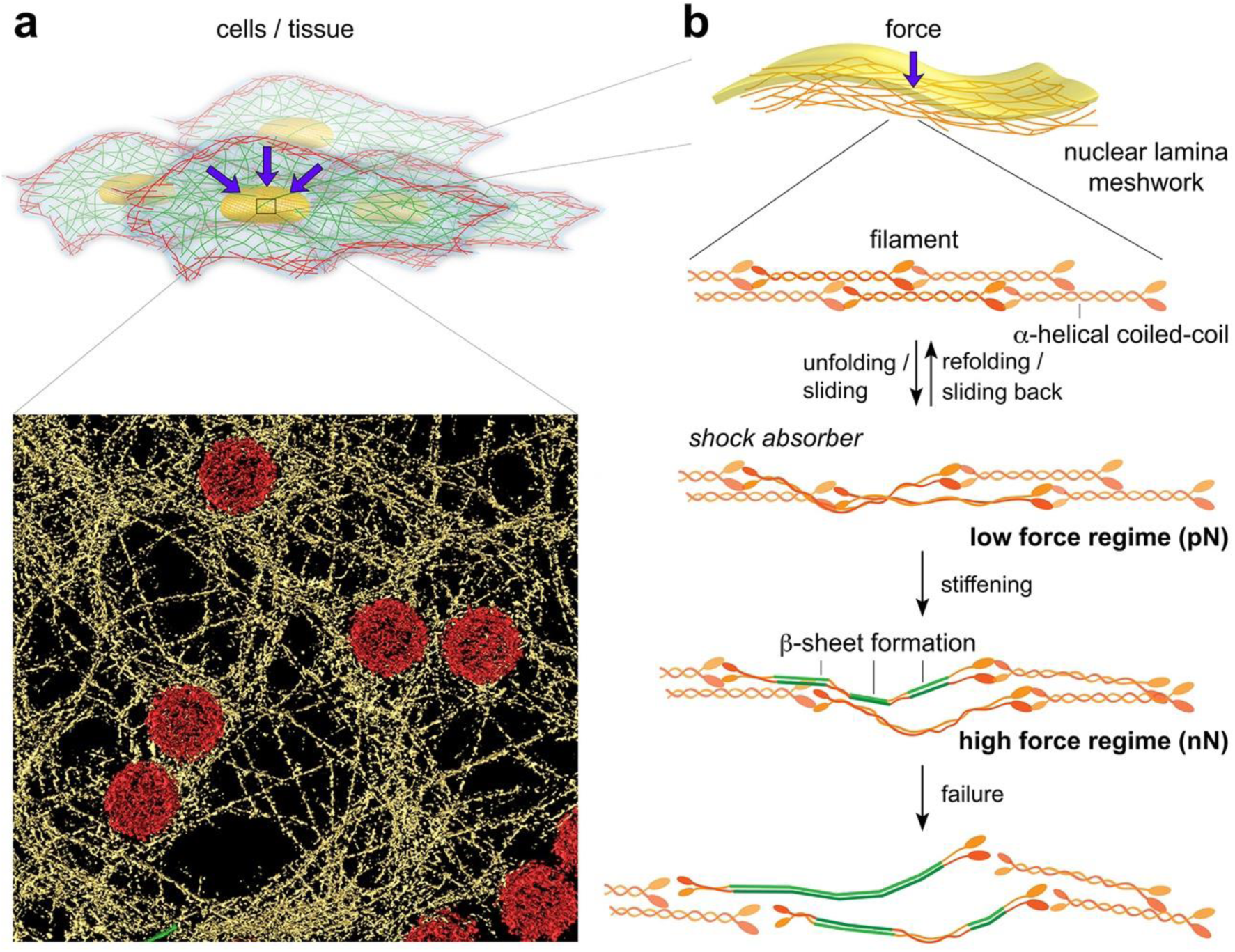
Nuclear lamins under external forces. (**a**) The cell nucleus is under constant stress from its surroundings and experiences continuous or prolonged mechanical shocks during division and migration^80, 81^. The nuclear lamina forms a meshwork that protects the genome and maintains the integrity of the nucleus. Surface rendered view of the lamin meshwork in MEFs. (**b**) A schematic model showing the response of lamin filaments when subjected to different levels of external force. Purple arrows denote point loads exerted on a nucleus, for example, by cytoskeletal elements^80^.

Interestingly, the strain propagation along the filaments in a lamin meshwork and the strength of lamin filaments up to failure increased with the meshwork connectivity. When λ = 0.5, increasing filament deformation caused a localized change in the meshwork strain and filament failure occurred at smaller strains (**Fig. 6a, b**). When λ = 5.6, increasing filament deformation led to a larger strain propagation in the meshwork peeling off the filaments from the underlying meshwork followed by local filament failure at higher strains (**Fig. 6c, d**). Furthermore, in a highly connected meshwork, filaments underwent larger deformation and were able to sustain higher forces as compared to a less connected meshwork (Supplementary Fig. 18). These results suggest that the lamin meshwork is an emergent structure; that is, the meshwork is more than the sum of its parts. The force required to damage an entire lamin meshwork by an outward pressure was determined to be ≈300 nN^70^, and the work done on the meshwork was estimated to be ≈10^8^ *k*_B_T. These high values on the meshwork compared to single filaments (≈10^5^ *k*_B_T) also point to the emergent nature of the meshwork which may be a key design feature.

### Mechanics of mammalian nuclei

Nuclear mechanics is known to be altered by lamin concentrations^71^. Our MD simulations and previous work^72^ suggest that the lamin meshwork topology determines filament and nuclear mechanics. To test this, we estimated the resistance or counter-force of isolated MEF nuclei by confining them between two parallel surfaces – a flat-wedged AFM cantilever and a glass surface^73^ (Supplementary Fig. 19). We observed that, (i) the counter-force generated by the nucleus on the cantilever increased as the confining space decreased; (ii) nuclei with either lamin A (lamin B knock-out) or lamin B (lamin A knock-out) alone showed higher counter-force than wild-type nuclei (Supplementary Fig. 20). The results suggest that nuclear mechanics depends on the concentration of the major lamin in the nucleus that also is likely to influence the meshwork topology.

## Discussion

Mechanical measurements of isolated nuclei^6, 25, 67^, intact cells^23^ or entire organisms^27^ by micropiperte aspiration, AFM and stretchable substrates provide insight into nuclear stiffness and morphology^74^. Such studies have assessed the changes to differing levels of lamins and NE proteins, and the underlying alterations of the physical properties^21^. Here, we characterized the mechanics of *in situ* assembled lamin filaments and meshworks providing a closer-to-native view of the physical properties of nuclear lamins. The experimental system allowed us to dissect the properties of lamin filaments from that of chromatin and other nucleoplasm components. Thus, we have bridged the gap between *in vitro* and *in vivo* by measuring filaments in the native lamin meshwork.

Combining mechanical and structural tools for interrogating individual lamin filaments offers a glimpse into the molecular mechanisms responsible for the mechanical properties of the lamin meshwork. The direct mechanical measurements of lamins *in situ* and meshwork simulations provide a mechanistic model into their role in protecting the nuclear contents in response to external forces (**Fig. 7**). The nonlinear behavior of lamin filaments under applied load and their connectivity in the meshwork confers exceptional strength and strain to the filaments. The filaments resist mechanical load by their stretching capacity rather than immediate breaking.

Mechanisms protecting the lamin meshwork against mechanical force are integrated at each level in the hierarchical construction of the meshwork – starting from the basic building block α-helix coiled-coil up to the higher order meshwork (**Fig. 7b**). The reversible unfolding or even sliding of the α-helical coiled-coil (rod domain) at low forces is the first protective step buffering mechanical shock given to a nucleus ensuring the structural integrity of the lamina and the nuclear contents (**Fig. 7b, middle**). At high forces, an irreversible strain-induced stiffening increases the filament strength, presumably by α-helix to β-sheet transition, further fortifying the safety mechanism against a possible meshwork failure (**Fig. 7b, bottom**).

Interestingly, the strength (failure force), toughness (energy dissipated or stored), and the resilience of the filaments increased as additional hubs were introduced in a meshwork (increasing λ from 0.5 to 5.6) (**Fig. 6e, f**). Point mutations in lamin A transform the nucleus from a resilient to a fragile material, most likely because of changes in the lamin meshwork topology^69^. Our results correlate the lamin meshwork topology and mechanical properties of single filaments, and may explain how re-modelling of the lamin meshwork may modulate nuclear mechanical properties, mechanotransduction and gene regulation^75^.

Here we have used the interactome of a subset of the nuclear lamin meshwork to model and simulate the meshwork mechanics for a direct comparison with experiments. The combination of AFM and cryo-ET together with network analysis is a novel approach that opens possibilities to understand the structural basis of nuclear mechanics in health and disease^72^. The application of network theory can be applied to quantitatively correlate the organization of the nuclear lamina to its mechanical role in diseases and malfunction by debilitating mutations^69^ in cells and tissues^71^. The integrative approach is a promising step towards combining structural mechanics and visual proteomics of entire cells and cellular organelles for obtaining insights into the function of macromolecular assemblies in health and disease. The conclusions also have implications for engineering lamins for material applications similar to silk and Kevlar^®^, and rationale design of protein-based meshworks with advanced mechanical functions^76^. For example, protein engineering of lamins combined with 3D printing technologies could be an area to explore in the near future.

## Methods

### *Xenopus laevis* nuclear lamina preparation for AFM measurements

*X. laevis* oocytes at stage VI were allowed to swell in a low salt buffer (LSB) (10 mM HEPES, 1 mM KCl, 1 mM MgCl_2_, pH 7.4) for 20 – 25 min. A prick with a sharp needle punctured the oocyte and enabled the nucleus to slowly squeeze out. The intact nuclei were immediately transferred to Modified Barth’s Buffer (MBB) (7.5 mM HEPES, 88 mM NaCl, 1 mM KCl, 0.4 mM CaCl_2_, 0.8 mM MgSO_4_, 2.5 mM NaHCO_3_, 2 mM Ca(NO_3_)_2_, TRIS to pH 7.5) and washed gently by a stream of the surrounding buffer repeatedly. The nuclei were then transferred to another Petri dish (World Precision Instruments) coated with poly-L-lysine (1 mg mL^−1^) which enabled the nuclei to stick firmly onto the glass surface of the dish. With the help of a glass microneedle the nucleus was slightly pushed onto the surface while rolling the needle to break open the nuclear membrane such that the nucleoplasmic side was facing upward, *i.e.*, inner nuclear membrane (INM). The nuclear contents including chromatin were gently removed, and the stuck nuclear membrane washed with ample volume of MBB (10 – 15 mL). For the experiment with Benzonase^®^ nuclease (Merck), the open nuclear membrane was incubated with 2500 U mL^−1^ of the nuclease for 1 – 2 h at room temperature.

### HeLa and mouse embryonic fibroblasts (MEFs) nuclear lamina preparation

Nuclear lamina of HeLa Kyoto and MEFs were prepared for imaging with AFM by unroofing the nuclei. Cells were seeded on autoclaved coverslips (#1 or 1.5, Carl Roth) and allowed to grow at 37 °C (5% v/v CO_2_) until a confluency of 75 – 90 % was obtained. The cells were prepared by washing the coverslips first with Ringer’s solution (+2 mM CaCl_2_) followed by Ringer’s solution without CaCl_2_. The coverslips were then exposed to hypotonic Ringer’s solution (one part of calcium-free Ringer’s solution was diluted in two parts deionized water) to swell the cells and facilitate easy opening^77^. To open the nuclei, a two-step procedure was developed. In the first step, cells were opened by placing an Alcian blue-coated coverslip on the cell-coated coverslip for ∼1 min. After ∼30 s, the excess buffer between the coverslips was wicked using a filter paper. After a further ∼30 s, the coverslips were separated by a stream of phosphate-buffered saline (PBS, pH 7.4) (500 – 1000 μL) using a pipette. Half-open cells with intact nuclei were transferred onto the top coverslip (Alcian blue-coated). The coverslips were then transferred to deionized water to swell the nuclei. The nuclei were then treated with Benzonase^®^ nuclease (Merck) in PBS (supplemented with 2 mM Mg^2+^) to digest the chromatin, washed with a high-salt buffer (PBS with 300 mM NaCl, pH 7.4), and then re-equilibrated in PBS. In the second step, the nuclei were opened to expose the lamina. For this, again an Alcian blue-coated coverslip was placed on the coverslip with nuclei, the excess buffer removed using a filter paper, and the coverslips separated by a stream of paraformaldehyde (4%). Both the coverslips were screened for nucleus and the one with higher density used for imaging. The procedure was carried out with and without protease inhibitors (Merck) without any noticeable effect on the lamin meshwork.

### AFM imaging and force spectroscopy

Nuclei were attached onto poly-L-lysine coated glass and were mechanically opened to ensure that the nucleoplasmic side was facing up, *i.e.*, accessible to the AFM cantilever (**Fig. 1a**). If the spread nuclear membrane folded (this happened usually at the edges because it could not stick to the poly-L-lysine surface), the sample were not measured in those regions. Since we always imaged and did our measurements in the center of the membrane, we are certain that all our measurements were conducted on the INM. In the AFM images of the lamin meshwork we took before pushing on the filament, no ruffles were observed. Moreover, folds and ruffles in the nuclear membrane can break the lamin meshwork.

Uncoated cantilevers of nominal tip radius ≍10 nm (HQ:CSC38/noAl, MikroMasch, Europe) were used for imaging and force spectroscopy. Before imaging and force spectroscopy, the cantilever sensitivity was determined by pressing the cantilever on a clean part (uncoated) of the glass surface. The spring constants measured using the in-built calibration module of the AFM (Nanowizard III, JPK instruments, Berlin) agreed with the typical range of the nominal spring constant 0.03 – 0.13 N m^−1^. *X. laevis* oocyte nuclei were imaged using quantitative force imaging with AFM. The samples were imaged at 128 × 128 pixels or 256 × 256 pixels. Random positions on lamin filaments were chosen in closed-loop mode (feedback on), and the tip of the AFM cantilever pushed on those positions with a force of 8 – 10 nN at different velocities (0.05 µm s^−1^ – 5 µm s^−1^). AFM experiments were performed in an acoustically isolated, temperature-controlled enclosure (26 ± 1 °C). Importantly, the data were collected over a span of >2 years providing reproducible results, ensuring that the quality and the selection procedure of the oocytes was maintained and did not affect the final results.

The force extension signals were exported in ASCII format from the JPK analysis software. The parameters: step unit (distance between the peak and the linear drop in the peak), failure force (peak of the force signal), stiffness (slope of the linear steep increase in the force signal), and deformation (distance between the inflection in the force extension signal and the first force peak) were all determined manually using Punias 3D. More than 50% of the data was analyzed 3 times to ensure the reproducibility of the manual procedure. The results were always in agreement within <5%. The loading rate was determined as the product of the empirically measured stiffness (slope of individual force signals) and the respective speeds (0.05, 0.1, 0.2, 0.5, 1, 2, 5 μm s^−1^).

In MEF somatic nuclei, the lamin meshwork is not in an orthogonal pattern (**Fig. 7**)^4^. Cryo-ET images from unswollen *X. laevis* oocyte nuclei also show that the native lamin meshwork is not always orthogonal (Supplementary Fig. 2). The elastic modulii (∼25 mN m^−1^) of lamin meshwork were estimated to be similar in swollen and unswollen nuclei of *X. laevis* oocyte showing that the lamin meshwork is capable of large changes while maintaining the material properties^5^. We are therefore confident that in our open nucleus system the meshwork is not perturbed to an extent that drastically changes filament mechanics and provides a close to *in situ* view of the meshwork.

### Reversible pushing of lamin filament

The sample preparation and positioning of AFM cantilever tip on lamin filaments was done as mentioned above. For the repetitive protocol, the cantilever was first pushed on chosen positions on lamin filaments with a force of 0.5 nN at a specific velocity (0.1 µm s^−1^ – 2 µm s^−1^) and then retracted 1 µm from the filament at a specific velocity (0.1 µm s^−1^ – 2 µm s^−1^). With a maximum retract velocity 2 µm s^−1^ of the cantilever over 1 µm, the refolding time of the α-helical coiled coil is estimated to be ∼500 ms. The cantilever was pushed again on the same spot with a specific velocity until the failure peak was detected.

### Nuclei purification for parallel plate assay

Cells were grown to 70 – 80% confluency in a T75 flask at 37 °C (5% CO_2_ v/v), washed with 10 mL PBS and treated with trypsin for 3 min at 37°C. Reaction was stopped by resuspending the cells in medium containing FCS; the suspension centrifuged at 1000 xg at 4 °C for 5 min. The cell pellet was washed with 5 mL cold (4°C) hypotonic buffer (10 mM HEPES, 1 mM KCl, 1.5 mM MgCl_2_.6H_2_O)^78^ and centrifuged at 1000 xg for 5 min at 4°C. The pellet was incubated in 5 mL hypotonic buffer containing 0.1% w/v digitonin on ice for 20 min, dounced 26 times, and centrifuged at 1000 xg for 5 min at 4°C. The pellet containing nuclei was resuspended in 5 mL cold hypotonic buffer, centrifuged (1000 xg for 5 min, 4°C), pellet resuspended in PBS (+2 mM MgCl_2_) and centrifuged again (1000 xg for 5 min, 4°C). The purified nuclei were re-suspended in 0.5 mL – 1 mL PBS (+2 mM MgCl_2_) containing 1% w/v BSA.

### Parallel plate assay for confining nucleus and measuring mechanical resistance

Focussed ion beam (FIB)-sculpted cantilevers were fixed on a standard JPK glass block and mounted in the AFM head (CellHesion 200; JPK Instruments). Nuclei were stained with NucBlue^®^ reagent (ThermoFischer Scientific). The bottom surface of the nucleus (attached to the glass) was focused and the cantilever approached to touch the glass surface close to the nucleus; this was set as 0 μm. The cantilever was retracted 20 μm – 25 μm away from the surface, positioned above the nucleus, and approached to touch the nucleus. Both, the piezo height and the force experienced by the cantilever, were monitored simultaneously in two different channels (Supplementary Fig. 19). Using the two signals and the point of cantilever deflection, the height of the nucleus was determined (Supplementary Fig. 20). Cantilever calibration was carried out using the thermal noise method (in-built calibration module).

### Cryo-electron tomography (cryo-ET) of nuclear lamina

*X. laevis* nuclear membranes were prepared in a similar manner as that for AFM experiments. Isolated nuclei were transferred onto a glow-discharged, perforated carbon copper grid (Quantifoil R2/1 200 mesh). The nuclei were opened manually and the nuclear envelopes spread over the grid. The grid was washed 3 times with LSB to remove residual chromatin and oocyte debris. A 3 μL drop of BSA-conjugated 10 nm colloidal gold was applied, and the grid was vitrified by rapid plunge-freezing in liquid ethane. Tilt-series (+60° – −60°) of the nuclear membrane was collected in a Titan Krios microscope (ThermoFischer Scientific) equipped with an energy filter and a K2 Summit direct electron detector using the dose fractionation mode at 5 fps. The projection images were acquired at a defocus of 6 µm at 42000x magnification, corresponding to a pixel size of 0.34 nm. The sample was exposed to a total dose of ≍ 52 e^−^ / Å^2^. The tomograms were reconstructed using TOM Toolbox and rendered in Amira software (v 6.0, FEI). Tomograms of MEFs nuclear lamina were acquired and rendered as described previously^4^.

### Meshwork analysis

Lamin filament density in individual slices of the tomograms were rendered manually using the Amira software. 13 tomograms from *X. laevis* nuclei and 12 from MEF nuclei were analyzed and rendered. The rendered 3D tomograms were automatically converted into graphs using Amira (Thermo Fisher Scientific). The vertices and links were then manually checked to verify the automated procedure. Closed loops, overlying links and extra vertices were manually deleted upon comparison with individual slices of the tomograms. The graph co-ordinates were then exported to Cytoscape (v 3.4.0) and the parameters analyzed using the network analyzer module.

### Molecular dynamics simulations

#### Scale free model of the lamin meshwork

The mesoscopic model of the lamin meshwork used here is based on a combination of experimental and full atomistic data. The geometry of the meshwork was obtained from our experimental observations that give its topological feature to be scale free. The meshwork was initially generated by deciding the connectivity of the end nodes of all the lamin filaments with the node list and their connectivities built according to a Barabasi–Albert model. This connected node list was further modified by repeating a simple Monte-Carlo process for 1,000,000 times to randomly select a node and change its connectivity. This change will be more likely to retain if it makes the degree probability distribution closer to the desired Power-law distribution (*P*(*k*) ∼ *k* ^−λ^) with the desired scaling factor λ. We randomly assigned the coordinates of all the nodes in a 2-dimensional plane that mimic the effect of nuclear membrane and treat each connection between the nodes as an existing lamin filament. Thereafter, we ran another simple Monte-Carlo process for 600,000 times and for each one we took the coordinates of a randomly selected node to have a random move. By doing so we ensured that the filaments in the meshwork have an average length of 12 ± 3 nm. The entire process was repeated to obtain meshwork models with different λ values. To account for the random effects introduced in the model generation, we performed individual loading tests on 100 meshworks and statistically studied the mechanical response from simulations.

#### Multiscale-modeling of lamin filament

We used mesoscopic beads to model each filament within the meshwork and fix *r*_0_ = 1 nm as the initial distance between neighboring beads as well as the equilibrium length of the springs. This length is much smaller than the persistence length of the lamin filament (≍ 1 um). We used a simple mesoscopic model describing each IF as a series of beads interacting according to nonlinear inter-particle multibody potentials that is obtained from full atomistic simulations^32, 51^. The total energy in our system is given by:

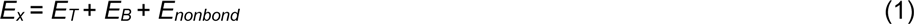

where *E_T_* is the energy term that associates with the tensile deformation of the IF fiber, *E_B_* is the energy term for the bending energy and *E_nonbond_* is the non-bonded energy term for two fibers in contact. We approximated the nonlinear force-extension behavior under tensile loading with a multi-polynomial potential model with its details and numerical parameters given below.

#### Computational experiments on the meshwork

Calculations were carried out in two steps, 1) relaxation followed by 2) loading. We modeled the effect of the nuclear membrane as a fixed plane substrate that has van der Waals interactions with the lamin filaments with a surface energy of 20 mJ m^−2^ at an equilibrium distance of 1.5 nm and cut-off distance of 4.0 nm. This energy is in the same order as vimentin protein adhesion on a silica surface and cell adhesion on a mineral surface^79^. Relaxation was achieved by heating up the system, then annealing the structure at a temperature of 300 K, followed by energy minimization. After relaxation, the system was maintained at 300 K in an NVT ensemble (constant temperature, constant volume, and constant number of particles) and loading applied by displacing a point within a single filament (by applying a point loading in the normal direction with the meshwork). While the single point was under a pushing force, the rest of the beads within the cut-off distance interact with the substrate, giving the reaction force to deform the filament. This set-up resembles the AFM pushing at a single point on a lamin filament, continuously displacing particles in the boundary at a speed of 0.1 mm s^−1^. It is confirmed that the loading rate chosen here is slow enough that leads to quasi-static deformation conditions.

## Reporting Summary

A reporting summary for this article is available as a Supplementary Information file.

## Data Availability

The data that support the findings of this study are available from the corresponding authors upon reasonable request.

## Acknowledgements

The authors thank Y. Turgay for discussions and providing tomograms of MEF nuclear lamina. KTS appreciates support from Forschungskredit fellowship, University of Zurich. This work was funded by a Swiss National Science Foundation Grant (SNSF 31003A_179418), the Mäxi Foundation to O.M. Additional support was provided by the Office of Naval Research (N00014-16-1-2333) to ZQ and MB. We thank the Center for Microscopy and Image Analysis at the University of Zurich.

## Author Contributions

KTS and OM conceived the concept of *in situ* lamin mechanics with input from DJM and UA. KTS developed the idea of combining mechanics and cryo-ET with MD simulations. ADG performed the cryo-ET experiments on frog oocyte nuclei. KTS performed the AFM experiments and analyzed the AFM and cryo-ET data. ZQ performed the simulations and analyzed the data. KTS wrote the paper with input from ZQ on MD simulations. All authors contributed to the final writing of the paper.

## Supplementary Data

### Note 1

#### Lamin filament mechanics under force

We first consider the deformation mechanics in the reversible low force regime, assigned to the unfolding of the α-helical coiled-coil domain^1^. A lamin filament ≈ 54 nm in length (*l*) (considering only the rod domain, 352 amino acids) if loaded in the center normal to the filament axis, undergoes an initial deformation, *d*_low_ ≈ 54 nm. At an average plateau force, *F*_low_ ≈ 0.3 nN, the filament would bend *θ* ≈ 63° with respect to the long axis of the filament [*θ* = tan^−1^ (*d*_low_ / (0.5 *l*))]. This is in agreement with the value determined for nuclei of cells spread on nano-pillars of radius 100 nm showing a nuclear membrane deformation of *θ* ≈ 58° ^2^.

Thus, the lamin filament (radius, *r* ≈ 2 nm) under an engineering stress, σ_plateau_ ≈ 24 MPa, would be stretched to a length (*L*_low_) ≈61 nm (122 nm the entire length) [*L* = *d*_low_ / sin(*θ*)]. The engineering strain, *ε*_low_, in the low force regime is estimated to be ≈126% which is similar to that measured for a vimentin dimer by atomistic simulations^3^. Because the plateau is reversible, we estimate the Young’s modulus to be ≈19 MPa.

Next, we estimated the mechanical parameters in the high force regime of the FE curve where the filament stiffens. Similar to the plateau region, the loading is assumed at the center of the filament. Thus, at a breaking load, *F*_high_ ≈ 3 nN, the filament was deformed to *d*_high_ ≈ 91 nm, normal to the filament axis. At a calculated yield stress, *σ*_yield_ ≈ 239 MPa, the filament would further bend at an angle *θ* ≈ 73° relative to its long axis. Stretched to a length (*L*_high_) of ≈95 nm (190 nm the entire filament), the engineering strain, *ε*_high_, of the filament is ≈250%. In such a scenario, the force acting along the filament axis (*F*_A_) would be ≈3.1 nN, in agreement with *F*_high_ ≈ 3 nN (normal to the axis). Thus, the loading of the filament bonds is presumably independent of the directionality of the force acting on the filament, *i.e.*, whether the filament is bent or stretched. This phenomenon is also illustrated in the similarity of FE curves obtained by pushing (**Fig. 1, Supplementary Fig. 6**) or stretching (**Supplementary Fig. 7**)^1^ lamin filaments in a meshwork.

---

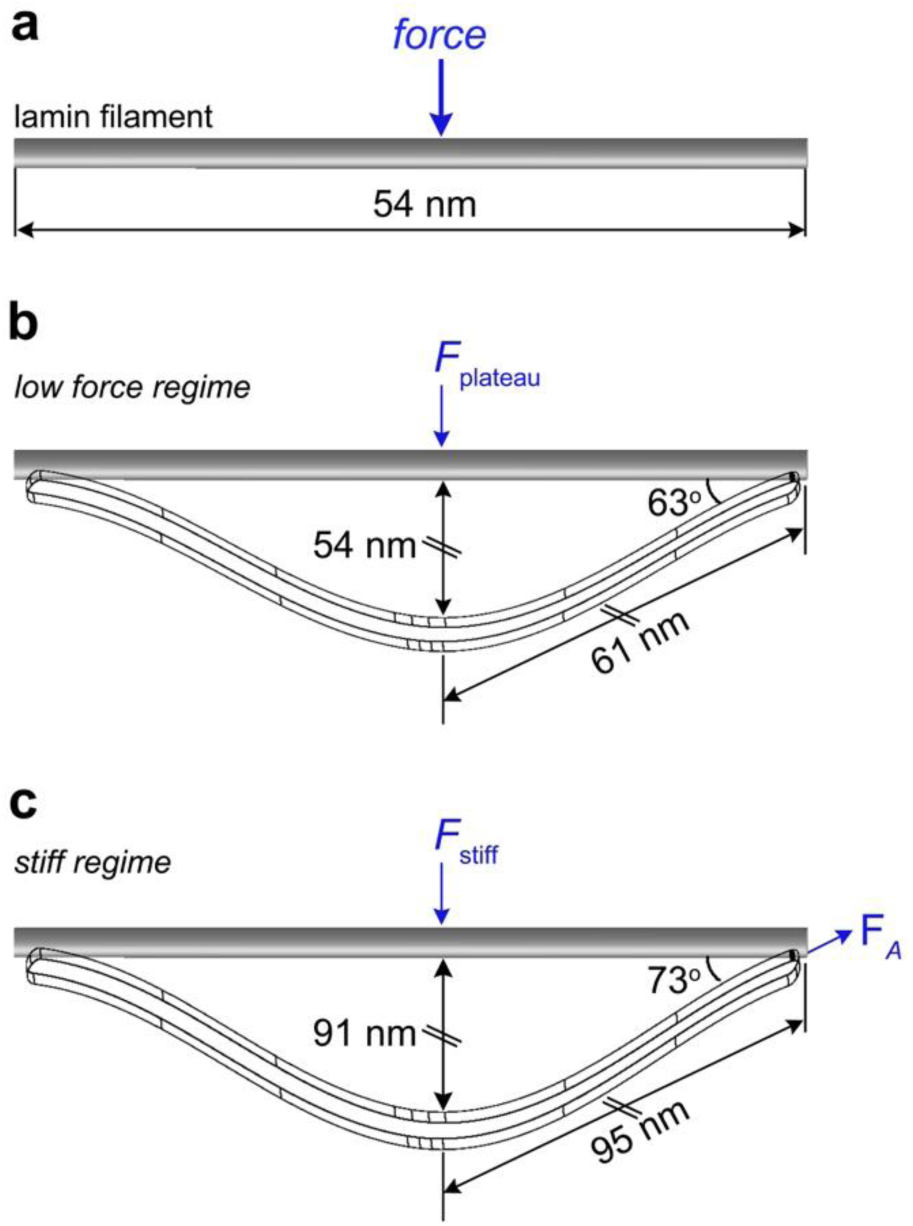
Model of the mechanical bending of a lamin filament. (**a**) Applying a force in the center of a 54 nm lamin filament results in a transformation through a reversible low force regime (**b**), followed by stiffening associated with α-helix to β-sheet transition (**c**).

**Supplementary Fig. 1.**
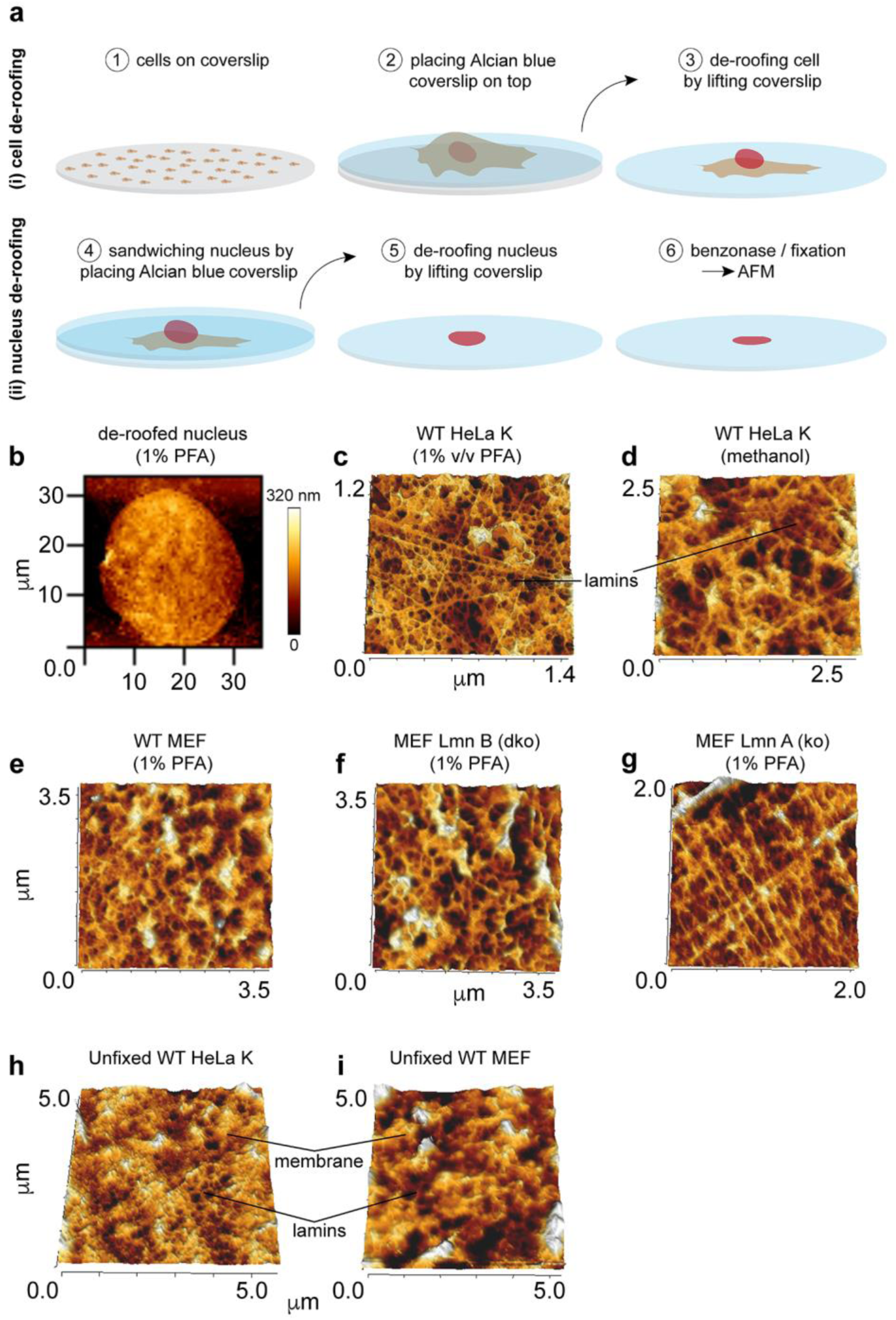
Imaging of mammalian nuclear lamina *in situ* by AFM. (**a**) A schematic description of the procedure used for imaging mammalian nuclear lamina with AFM. HeLa cells seeded on a coverslip were unroofed by placing and lifting an Alcian blue-coated coverslip. This facilitated the transfer of de-roofed cells with intact nuclei onto the Alcian blue coverslip. Nuclei were de-roofed in a second step by again placing and lifting an Alcian blue-coated coverslip. The open nuclei were chemically fixed and imaged with AFM showing filamentous structures, lacking NPCs. (**b**) A typical AFM topograph of a de-roofed nucleus obtained by quantitative imaging. (**c, d**) Images of the lamin meshwork from wild-type (WT) HeLa K cells, chemically fixed with 1% v/v PFA and methanol, respectively. (**e – g**) Images of lamin meshwork from wild-type, B-type lamin knockout and A-type lamin knockout MEF cells fixed with 1% v/v PFA^4^. Most likely the NPCs were removed during the de-roofing and therefore were not observed. (**h, i**) Nuclei that were not chemically fixed showed more membranes and not a clear meshwork as observed in fixed nuclei. Therefore, mechanical measurements on single lamin filaments were not performed on unfixed mammalian nuclei.

**Supplementary Fig. 2.**
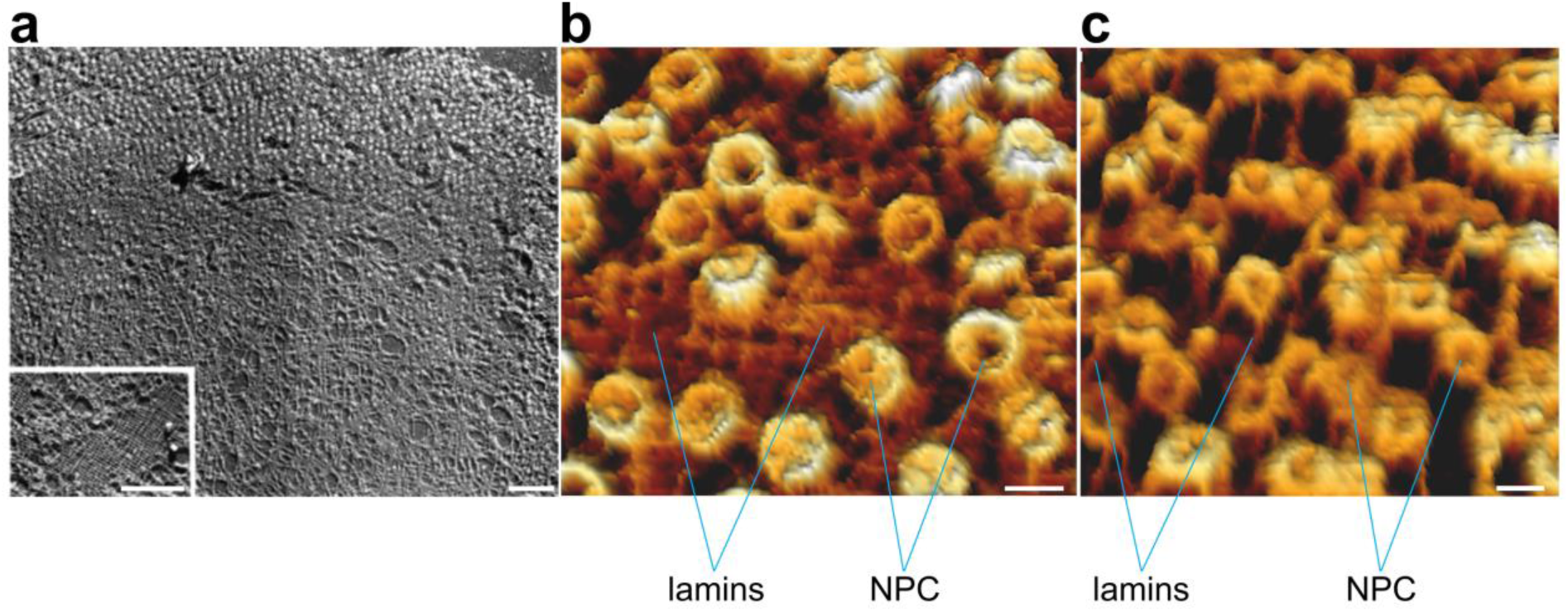
Electron microscopy and atomic force microscopy imaging of *X. laevis* oocyte nuclear envelopes (NEs). (**a**) Previously published image of a NE showing lamin meshwork and NPCs, obtained using detergent treated, freeze-dried / metal-shadowed NE from *X. laevis* oocyte nucleus^5^. The meshwork shows lamin filaments arranged orthogonally and in an apparently random manner. Image used with permission from ref. *5*. Scale bars, 1000 nm. (**b, c**) Images of NEs obtained in this study from nuclei of *X. laevis* oocytes that were isolated and manually opened to expose the nucleoplasmic face of the organelle. The orthogonal arrangement of the lamin filaments in the meshwork (and the NPCs) was clearly resolved by quantitative force imaging with AFM. Other regions of the samples showed less ordered lamin filament organization at the NE. Scale bars, 100 nm.

**Supplementary Fig. 3.**
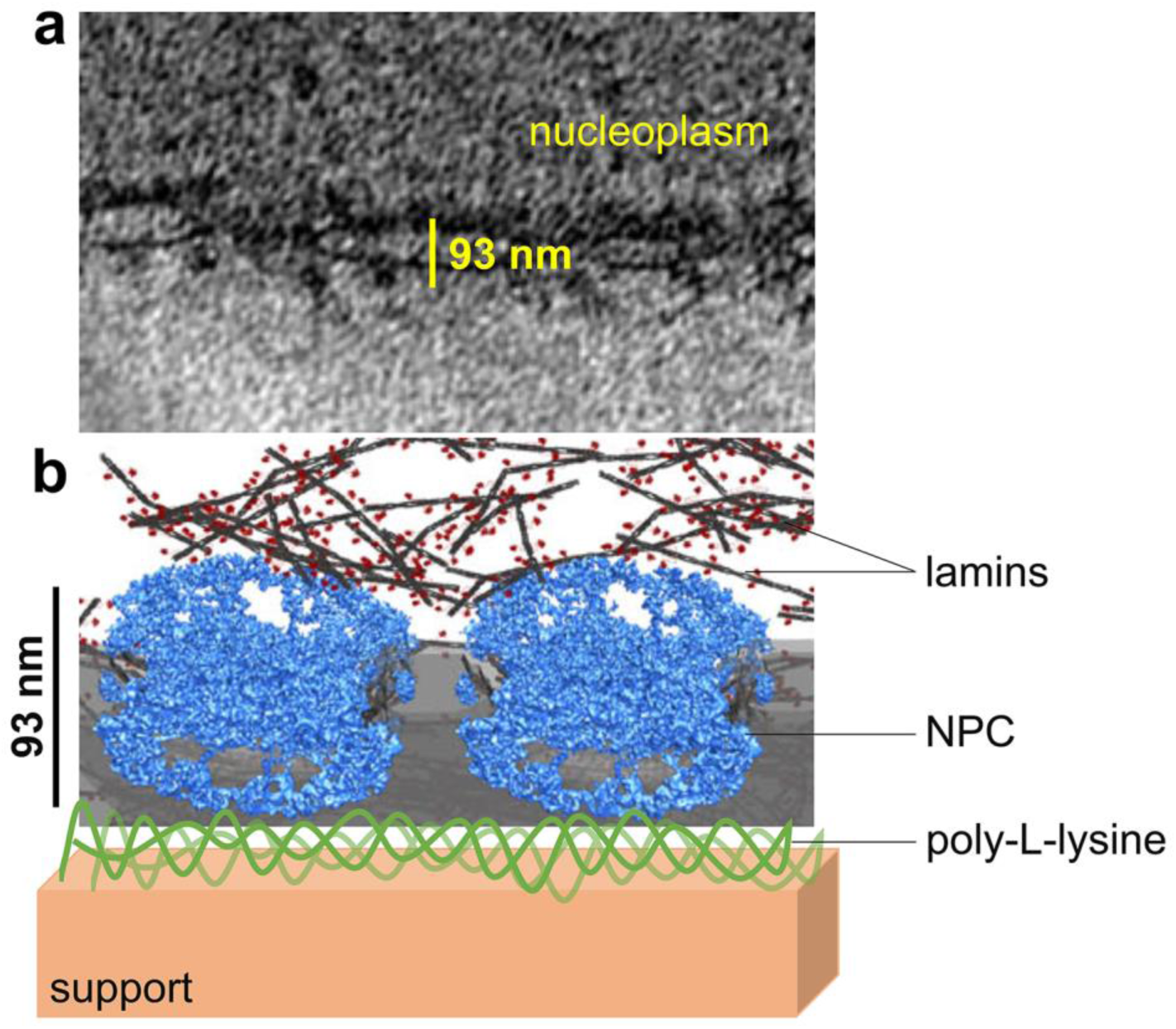
Lamin meshwork is situated at a distance of >93 nm from the surface support. (**a**) An *x-y* slice through a tomogram showing the distance between the INM and the cytoplasmic densities at the ONM, at the NE of a *X. laevis* oocyte (adapted from ref. *6*). (**b**) A schematic view of our experimental setup. The glass surface was coated with poly-L-lysine. On top of the coated glass surface, the nuclear envelope was spread. Therefore, the lamin filaments and the glass support were separated by ≈100 nm.

**Supplementary Fig. 4.**
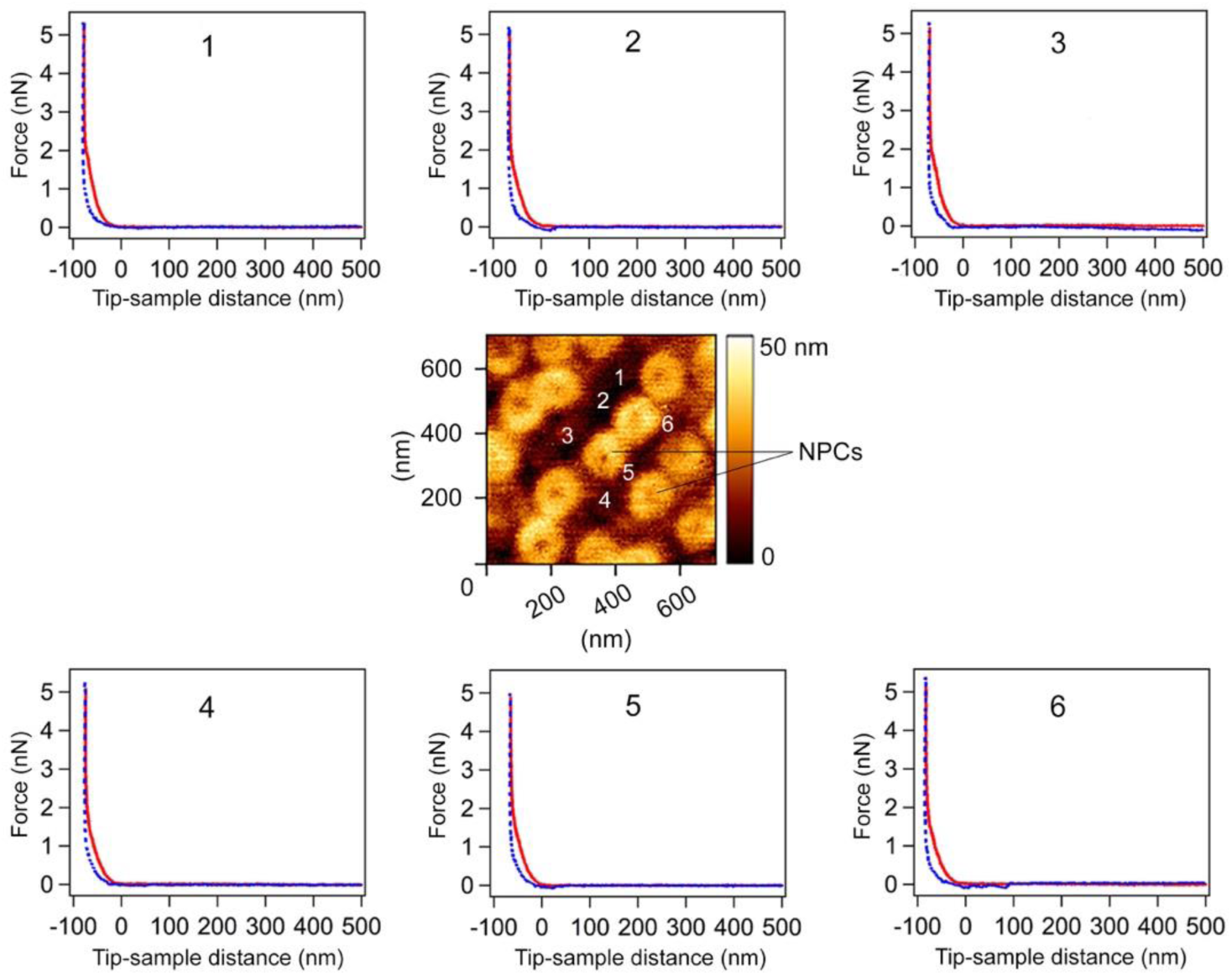
FE curves on nuclear membrane do not show force peaks. An AFM topograph was recorded by FE curve imaging. The region showed only NPCs but no lamin filaments. Positions marked 1-6 were chosen in between the NPCs and the cantilever tip pushed in the active feedback mode. No distinct force peaks were observed in the FE curves as observed in the curves obtained by pushing lamin filaments (Supplementary Figs. 5, 9). Red curves denote the approach (pushing) of the cantilever tip, blue curves denote the retract. The numbers (1-6) on the AFM topograph correspond to the numbers labelling the FE curves.

**Supplementary Fig. 5.**
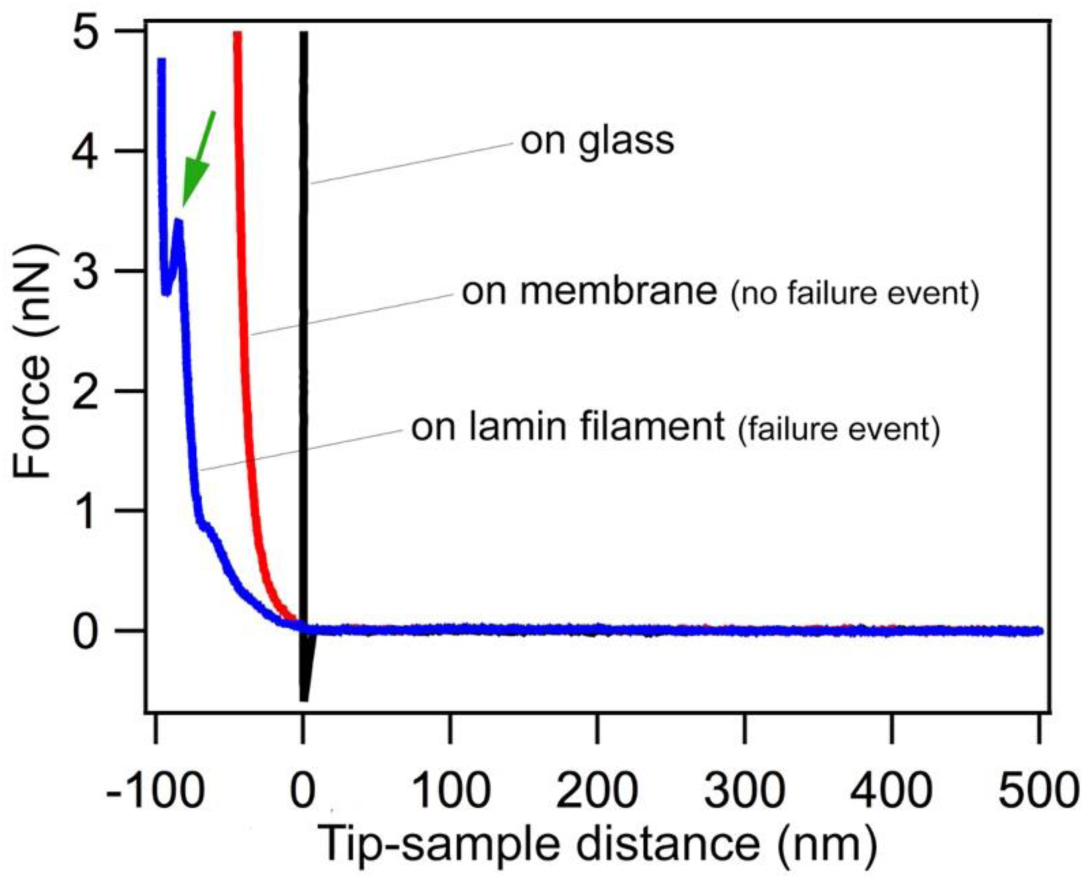
FE curves on glass, on membrane and on lamin filament show different profiles. The blue FE curve obtained by pushing a lamin filament shows an apparent failure event between 3 nN and 4 nN. On nuclear membrane (red FE curve) and bare glass without poly-L-lysine coating (black FE curve), no force peaks were observed. The FE curves clearly show that the surfaces are of different stiffness.

**Supplementary Fig. 6.**
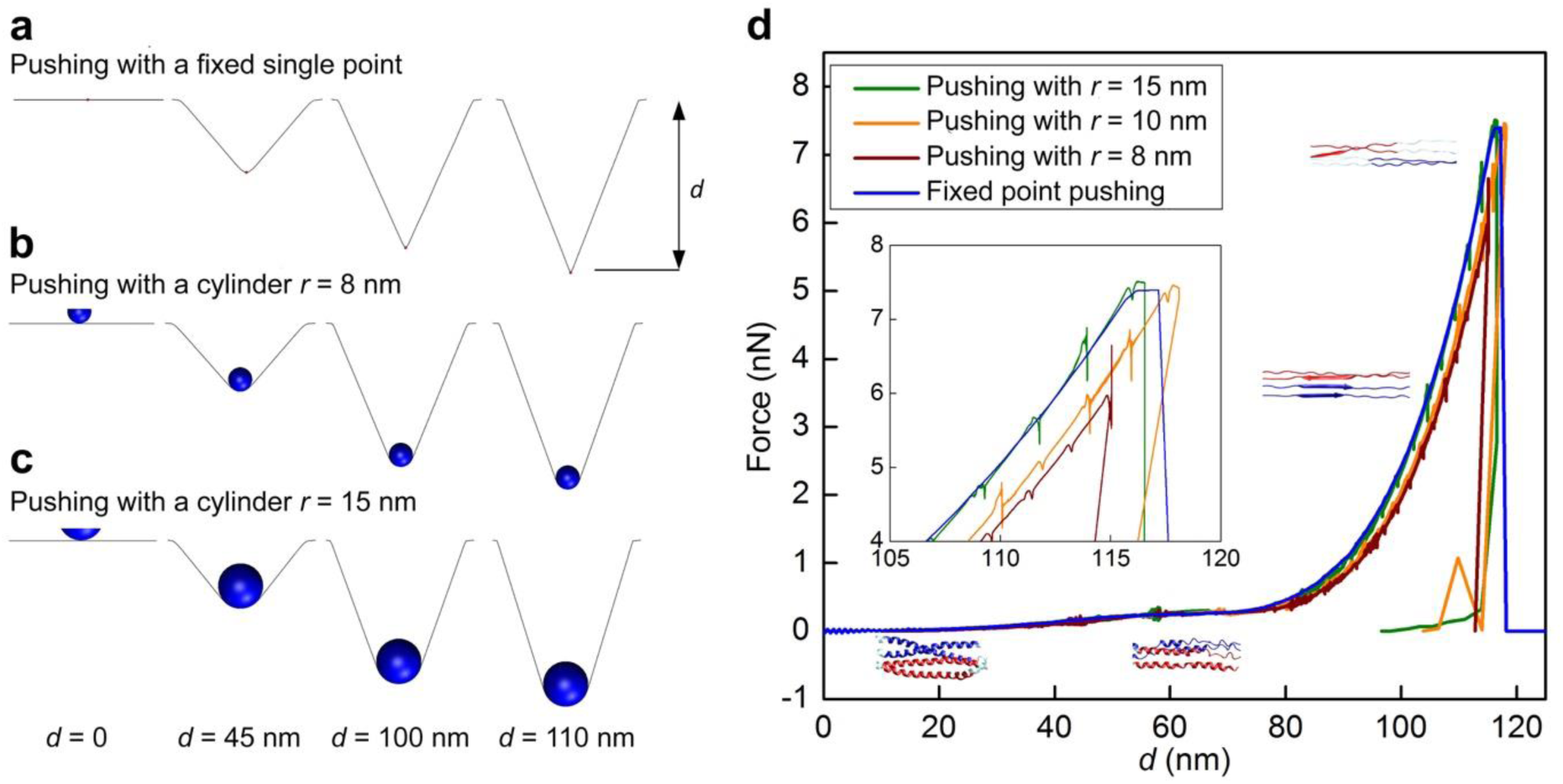
Lamin filament mechanics is not sensitive to cantilever tip curvature. (**a – c**) Simulation analysis confirmed that pushing with spheres of finite radii (AFM tip) is equivalent to pushing with a single fixed point. The AFM cantilever tip is represented by a sphere of finite radius (*r*). The tip is pushed in the middle of a single lamin filament of 100 nm length and with two fixed ends. The deformation snapshots at different displacement and the full force-displacement curves are given for the tips with different radii. For the pushing test with a fixed single point, only the middle of filament is subjected to the pushing force. (**d**) The different tests provided very similar results for the FE curves demonstrating that the loading can be simplified without modeling the contact between the cantilever tip and the filament. The structural states during the pushing of a lamin filament are shown at corresponding positions along the FE curves.

**Supplementary Fig. 7.**
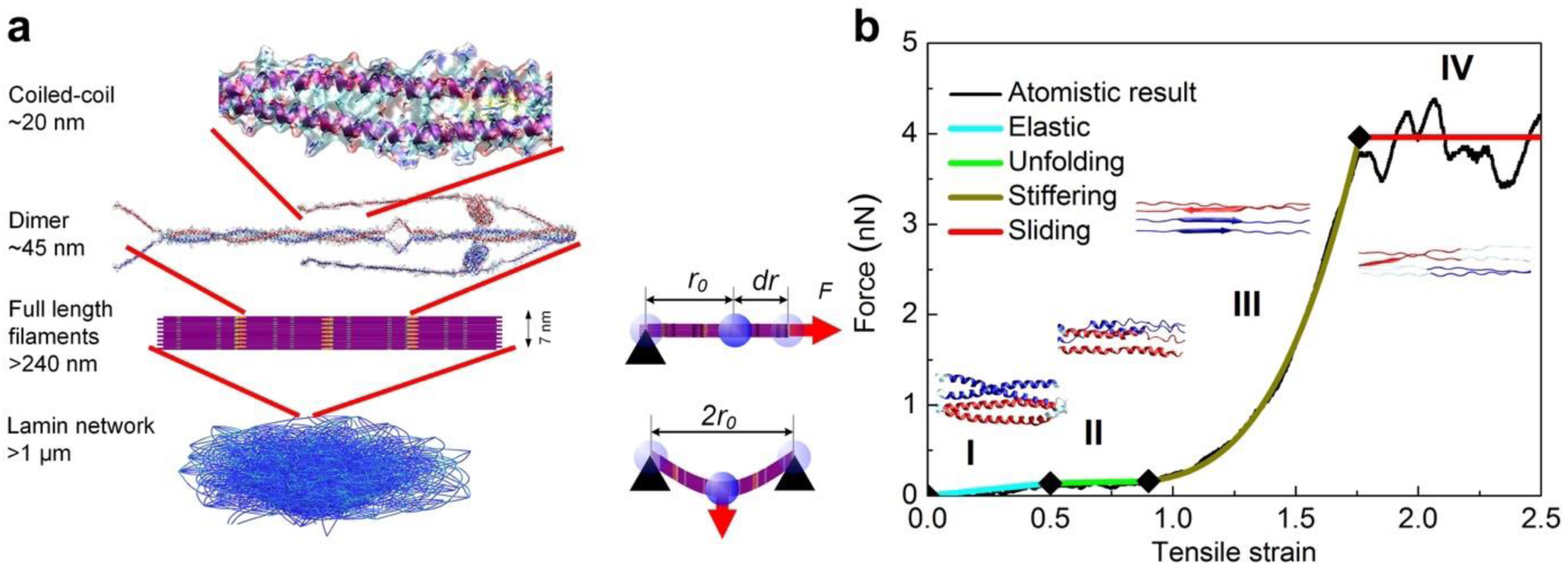
Stretching a lamin filament in the meshwork leads to similar structural transitions as pushing a filament. (**a**) Cartoon depicting the scale of meshwork to filaments to a dimer to an α-helical coiled-coil. (**b**) MD simulations of stretching a lamin filament along the horizontal plane of the lamin meshwork show (I) elastic stretching, (II) unfolding of α-helical coiled-coils, (III) stiffening because of α-helix to β-sheet transition, and (IV) sliding of the β-sheets leading to filament failure. Except the sliding of β-sheets at failure, the same structural states were observed when a lamin filament was pushed in the meshwork (Supplementary Fig. 6). Figure adapted from ref. ^1^.

**Supplementary Fig. 8.**
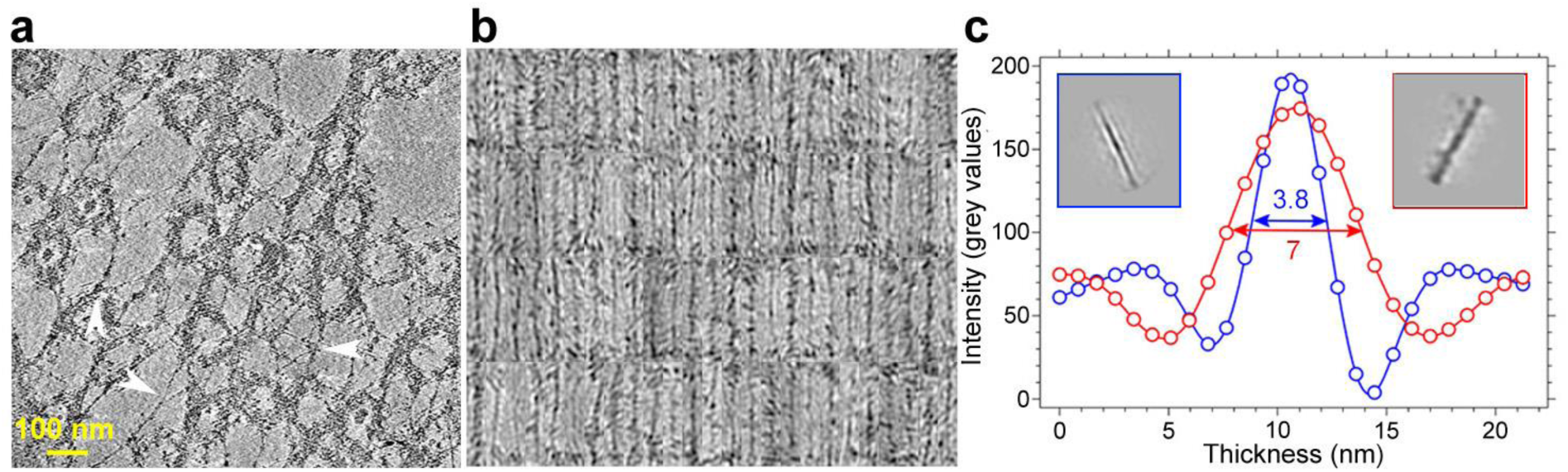
Lamin filaments analyzed by cryo-ET. (**a**) A 10 nm slice through a tomogram of spread NE showing the nuclear lamina of *X. laevis* oocyte. Lamin filaments often interact laterally to form thicker filaments (arrowheads). The filamentous lamin meshwork is anchored at NPCs (round densities). (**b**) A gallery of single lamin filaments extracted *in silico* from the tomograms. (**c**) Density lines through the rod-like structure of two structural class-averages (framed blue and red) indicating filament thickness of ≈3.8 nm and ≈7 nm in agreement with a tetramer and lateral association of tetramers, respectively. The filament dimensions are in agreement with previously measured nuclear lamin filaments in the MEF nuclei^7^. The structural class averages (the two prominent classes are shown in **c**, insets) were calculated as reported in ref. *7*. In brief, the filaments were identified and 1400 sub-volumes of 64 × 64 × 64 pixels (pixel size 1.36 nm, binning 2) were extracted. The filaments in these sub-volumes were then aligned to a cylindrical template parallel to the *y*-axis and projected in the *z*-direction. These projection images were then visually inspected with an interactive tool (available upon request) in MATLAB (Mathworks, Natick, USA) and subjected to 2D classification using Relion^8^.

**Supplementary Fig. 9.**
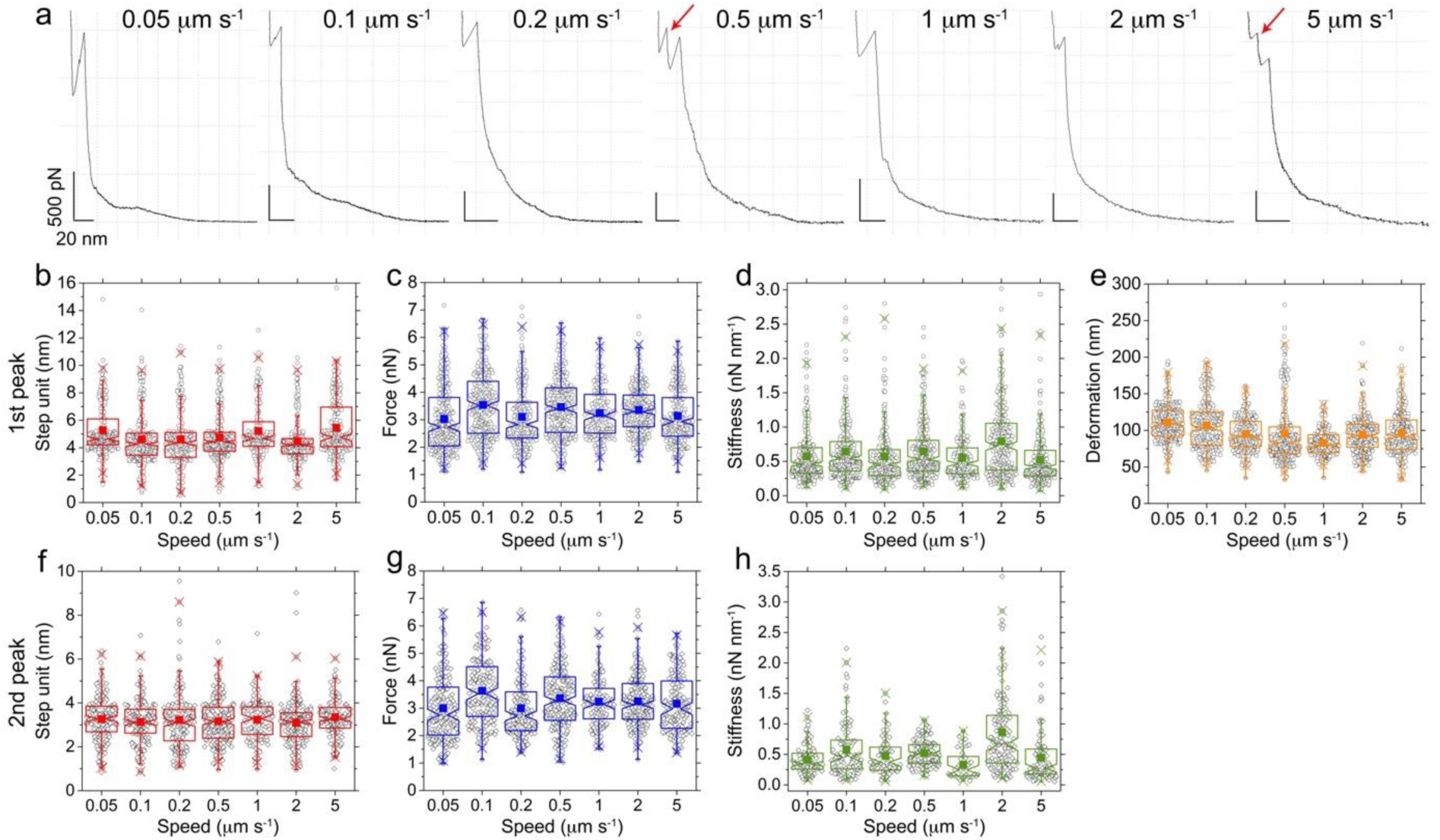
Notch box-plots of mechanical parameters at increasing loading rates. (**a**) Representative FE curves showing the mechanical behavior of lamin filaments under force. In some cases, after the first failure peak a second peak (red arrows) was observed. The FE curves were independent of the loading rates suggesting a similar load-bearing mechanism of lamin filaments at different loading rates. The step unit, force, stiffness and deformation of the first peak (**b – e**), and the second peak (**f – h**), did not change with the pushing speed. The box includes 25 – 75% of the data, each black circle being an event from a single filament. The solid squares denote the mean and the whiskers signify 1 – 99% of the data. The number of events analyzed at each speed for both the peaks is reported in Supplementary Table 2.

**Supplementary Fig. 10.**
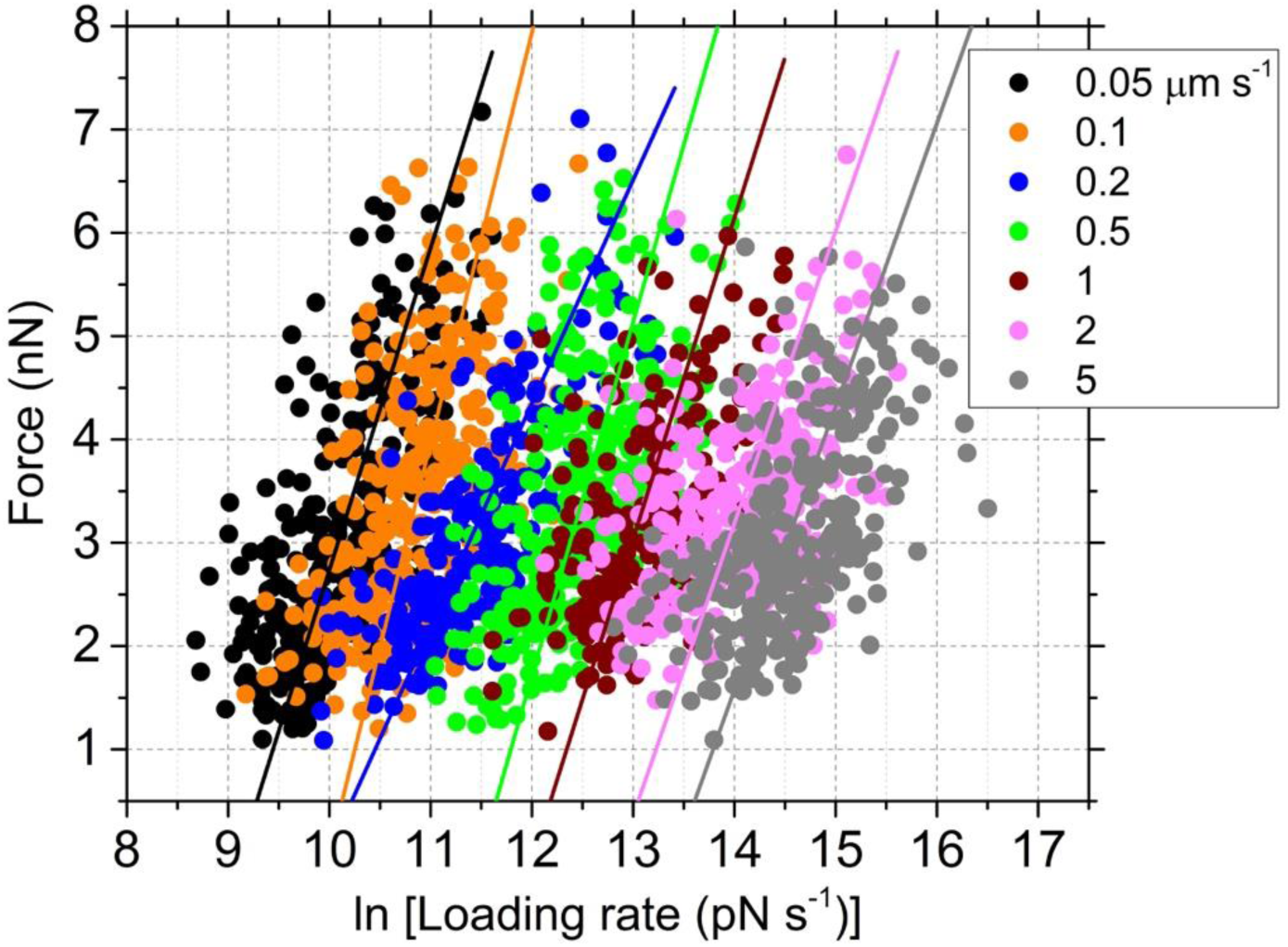
Apparent failure of lamin filaments is independent of loading rate. Each dot denotes a single force peak event detected in an FE curve. The number of events (*n*) at 0.05 µm s^−1^, *n* = 321; 0.1 µm s^−1^, *n* = 301; 0.2 µm s^−1^, *n* = 239; 0.5 µm s^−1^, *n* = 313; 1 µm s^−1^, *n* = 203; 2 µm s^−1^, *n* = 293; 5 µm s^−1^, *n* = 276. The failure forces of lamin were independent of the loading rate (stiffness × speed).

**Supplementary Fig. 11.**
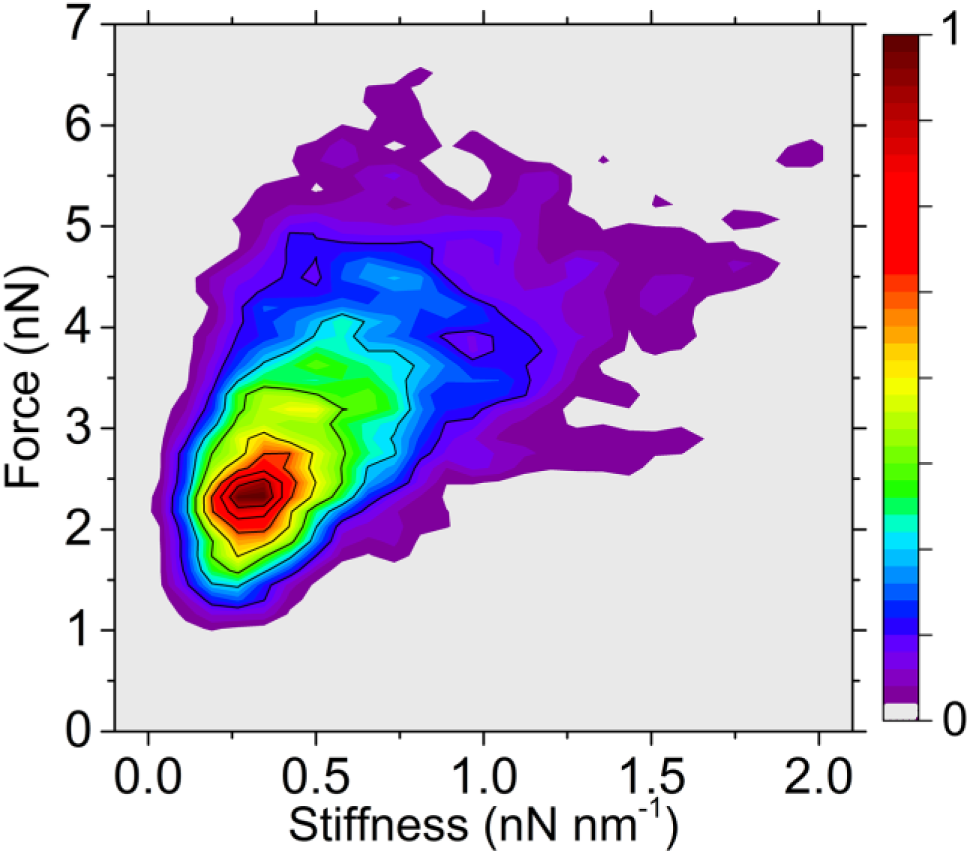
Correlation between force and stiffness of lamin filaments. Density map showing that the stiffer a filament becomes under force, it can tolerate higher forces before failure.

**Supplementary Fig. 12.**
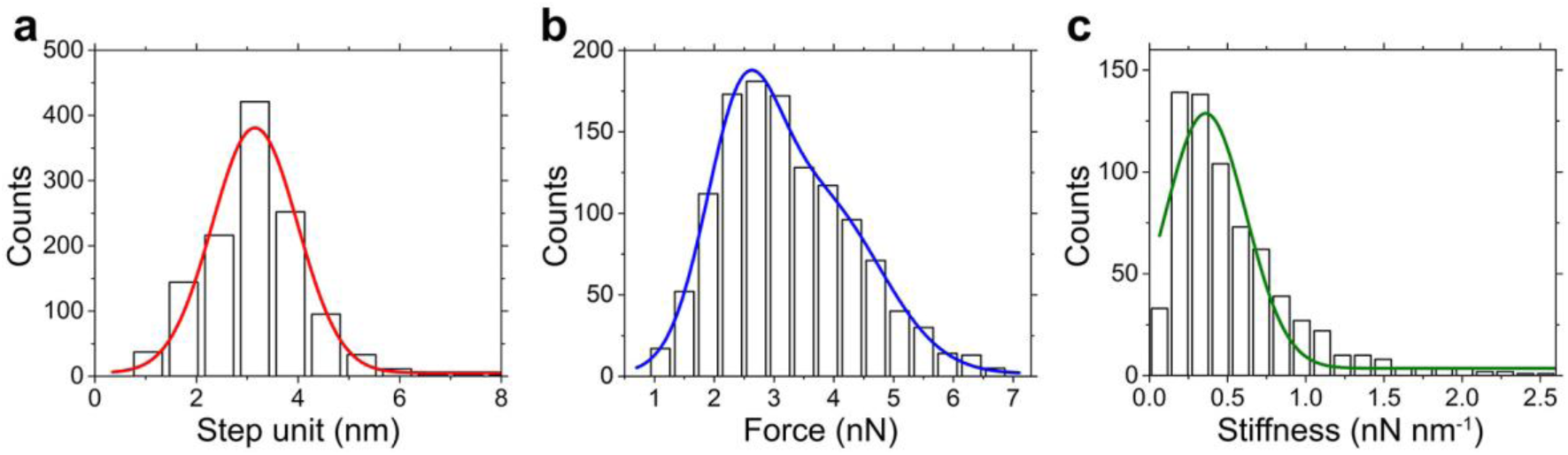
Analysis of the second peak in the FE curves of lamin filaments. Data from applying forces at different loading rates was pooled to plot histograms of (**a**) step unit (3.2 ± 0.9 nm, ± S.D., *n* = 1221), (**b**) failure force (2.5 ± 0.6 nN, 3.7 ± 1.0 nN, *n* = 1221), and (**c**) stiffness (0.36 ± 0.26 nN nm^−1^, *n* = 692), which were all similar to the values obtained for the first peak (Fig. 2).

**Supplementary Fig. 13.**
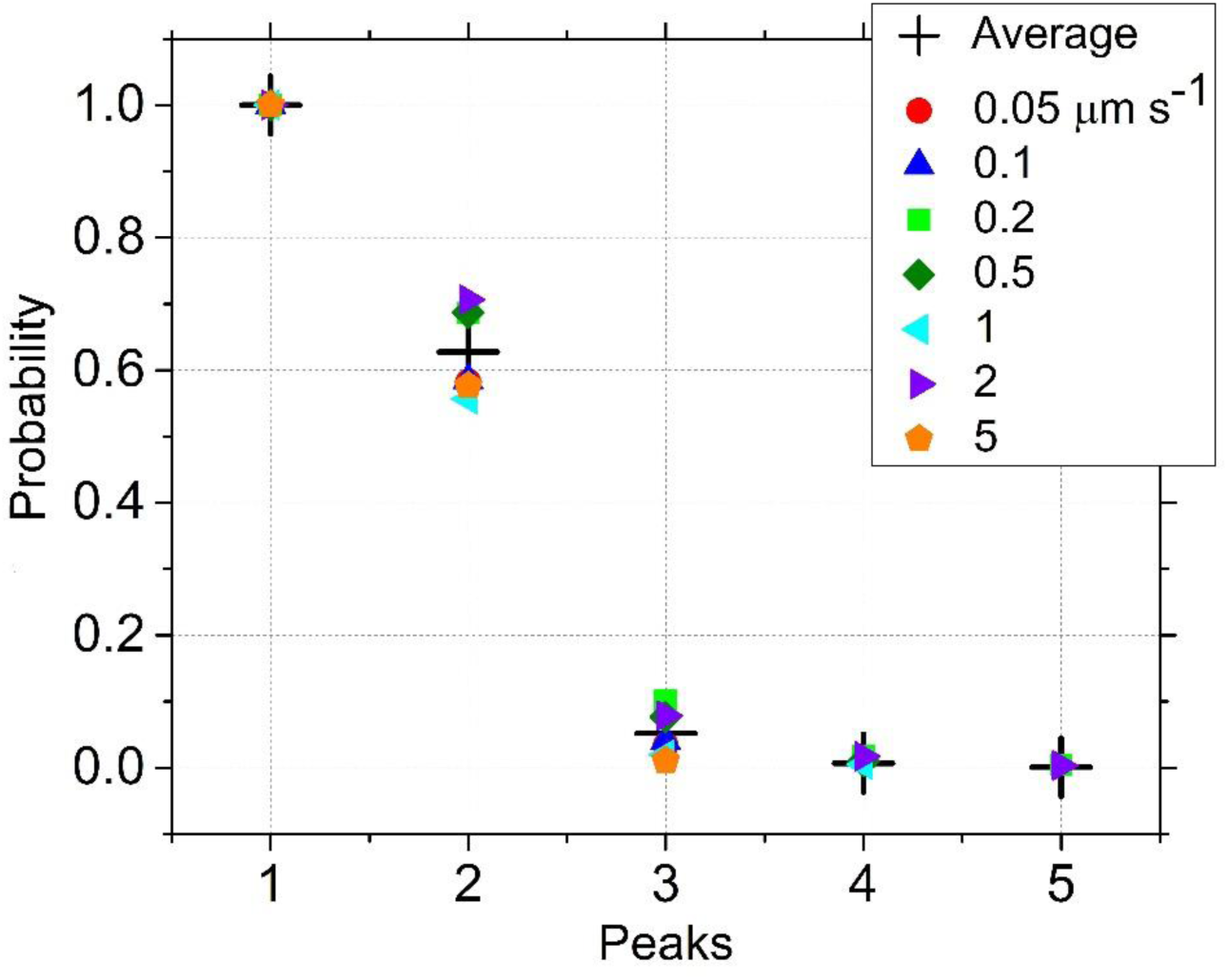
The detection probability of the mechanical failure of lamin filaments. The probabilities (normalized with respect to the first failure peak) measured in the FE curves of lamin filaments. Because the third, fourth and fifth peaks appeared rarely, only the first two peaks were considered for determining the step unit, force, stiffness and deformation. The values are reported in Supplementary Table 3.

**Supplementary Fig. 14.**
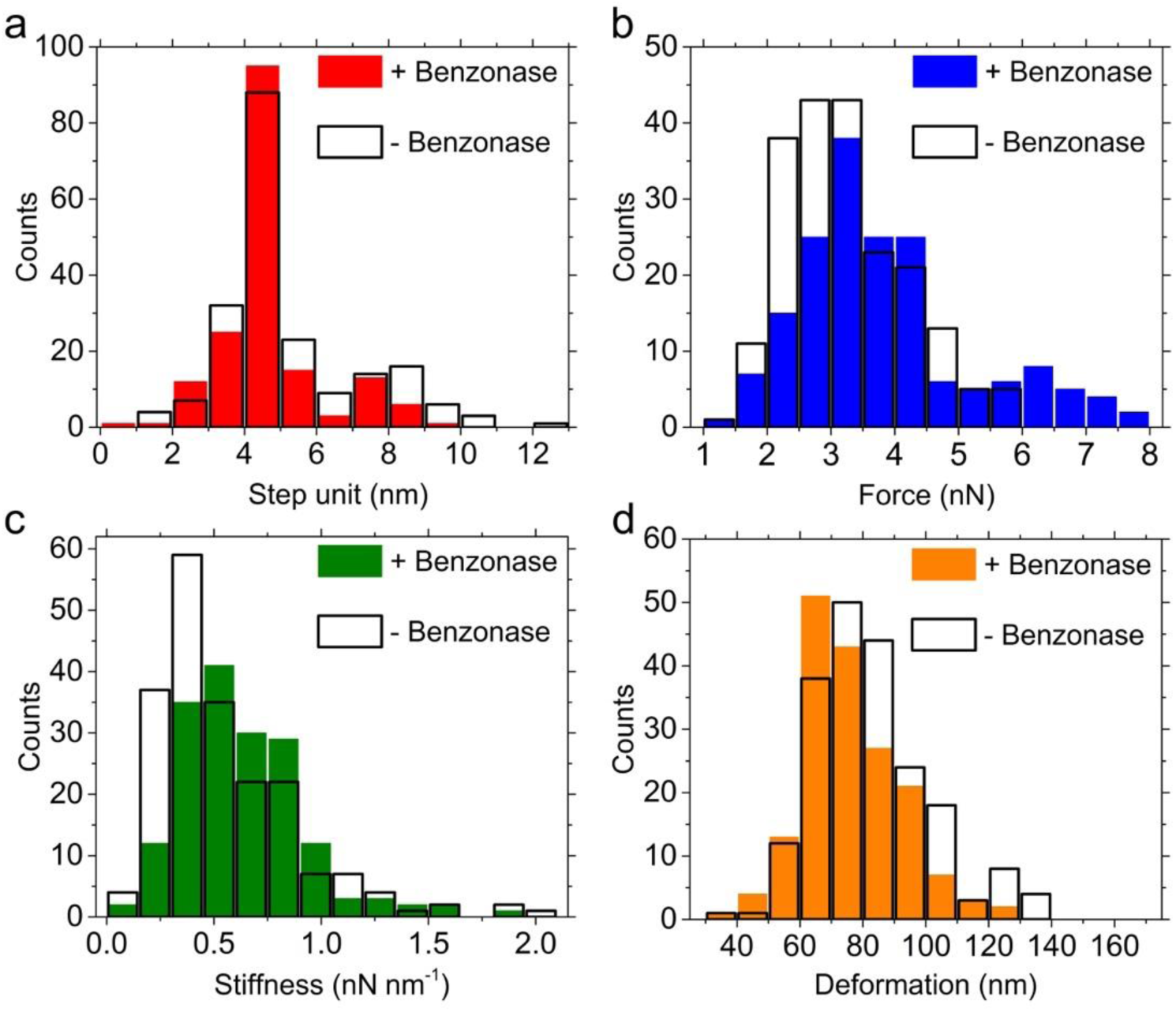
Benzonase nuclease treatment does not change lamin mechanics. (**a**) Step unit, (**b**) force, (**c**) stiffness and (**d**) deformation of lamin filaments in the meshwork of *X. laevis* nuclear envelopes were measured after nuclease treatment at a pushing speed of 1 µm s^−1^. The values of the parameters were comparable to those obtained without nuclease treatment, at 1 µm s^−1^, suggesting that chromatin, ribonucleoproteins or RNA were not present in the experimental system and did not influence the mechanical properties of lamins in our experiments. – Benzonase-free NEs, *n* = 204 for each parameter; + Benzonase-treated NEs, *n* = 172 for each parameter.

**Supplementary Fig. 15.**
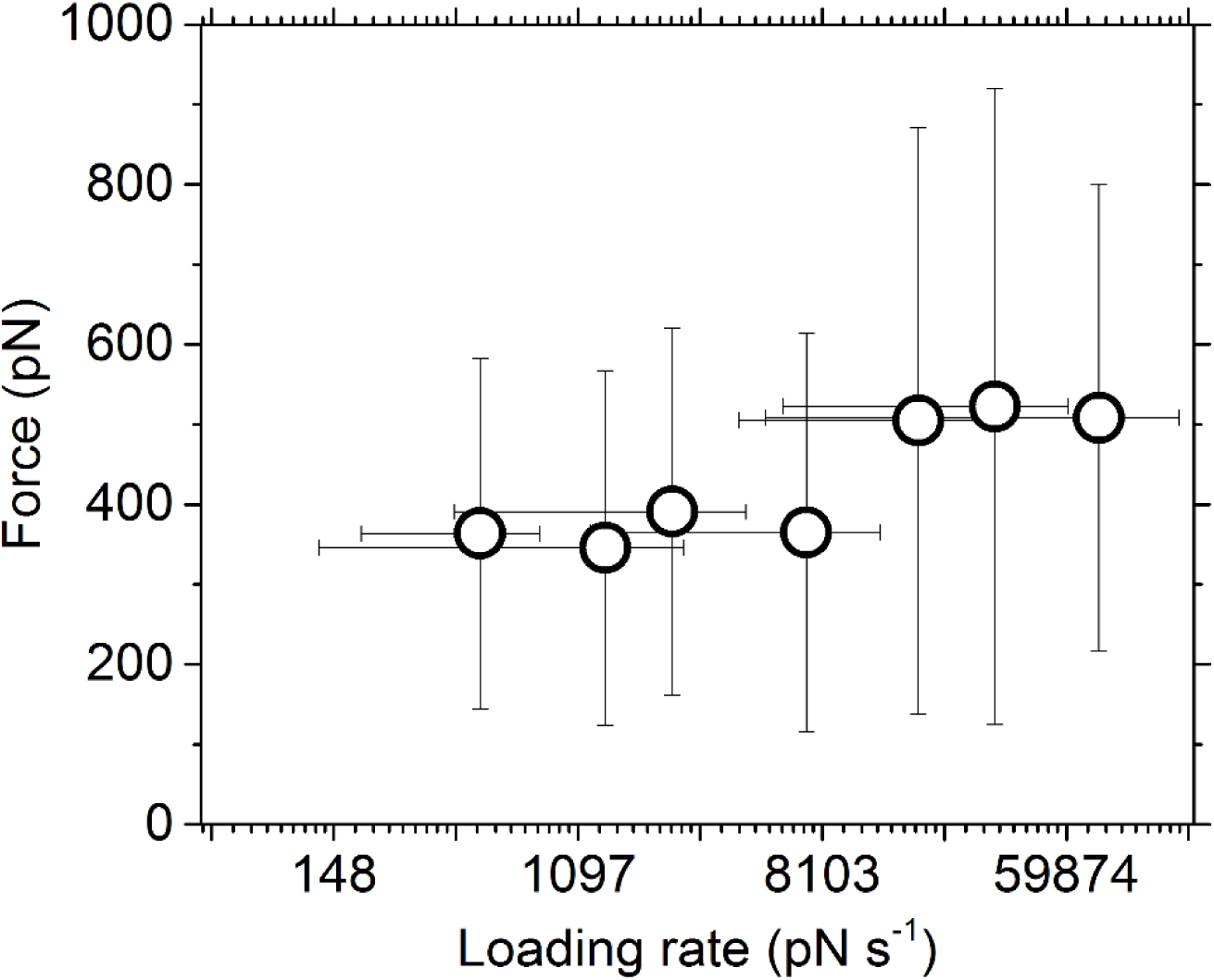
Mechanical response of lamin filaments at low force is independent of the loading rate. Similar to the failure force of the filament after stiffening, the force at which the plateau was observed in the low force regime did not change with increasing loading rates. The data points denote the average values ± standard deviations of the force (vertical error bars) and loading rate (horizontal error bars).

**Supplementary Fig. 16.**
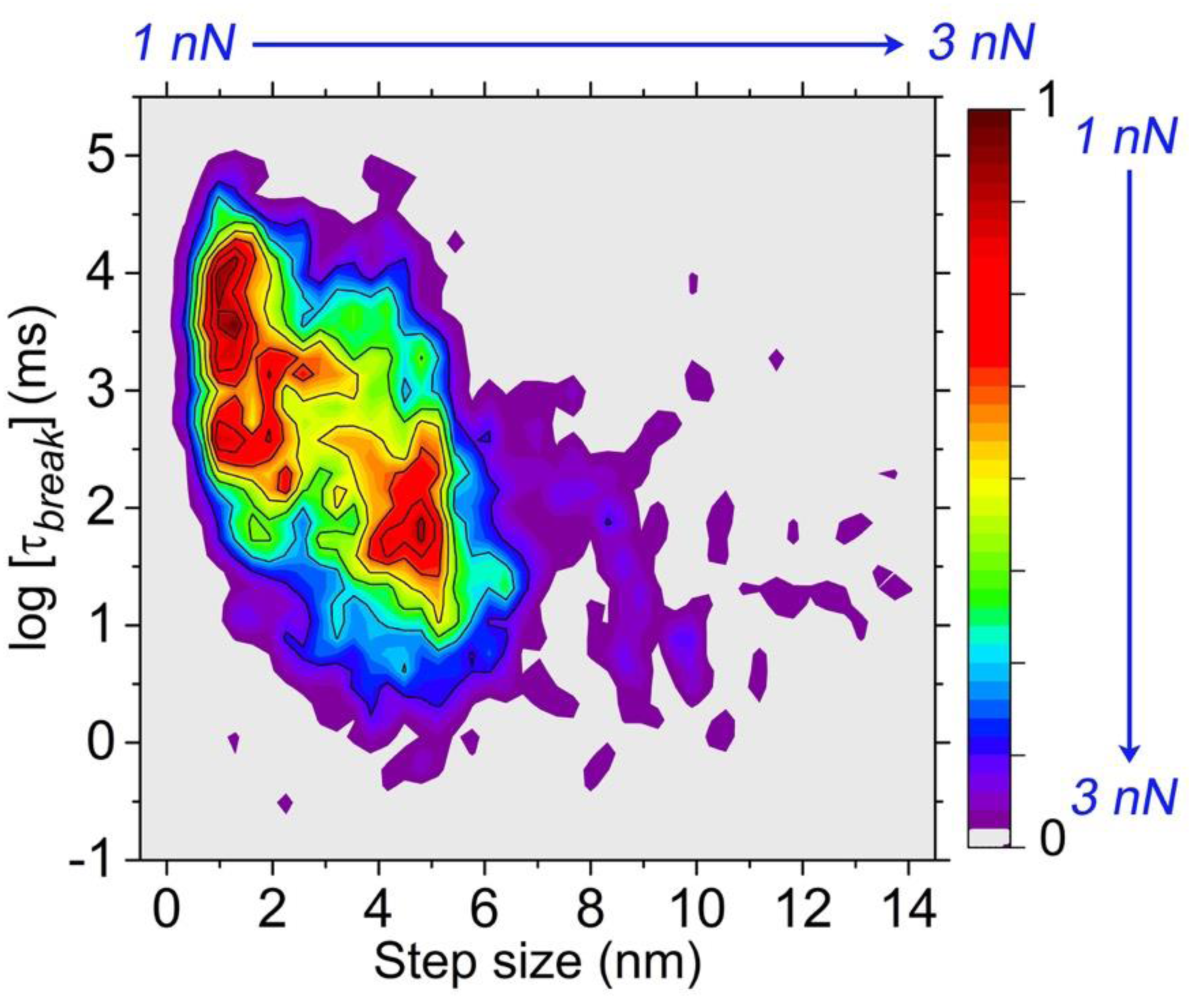
Lifetime of lamin filaments at constant loads. Density map showing the exponential decrease of the lifetime, τ*_break_*, with the step size (assigned to α-helix coiled-coil and β-sheet). Two dense populations (red areas) can be clearly seen in the plot. The axes (top and right) denote the trend of F*_load_*. α-helical elements (1 – 2 nm) can bear forces up to 1 nN for up to a hundred milliseconds, whereas tetramers (4 nm) stiffen at high forces (3 nN) and fail within a few milliseconds.

**Supplementary Fig. 17.**
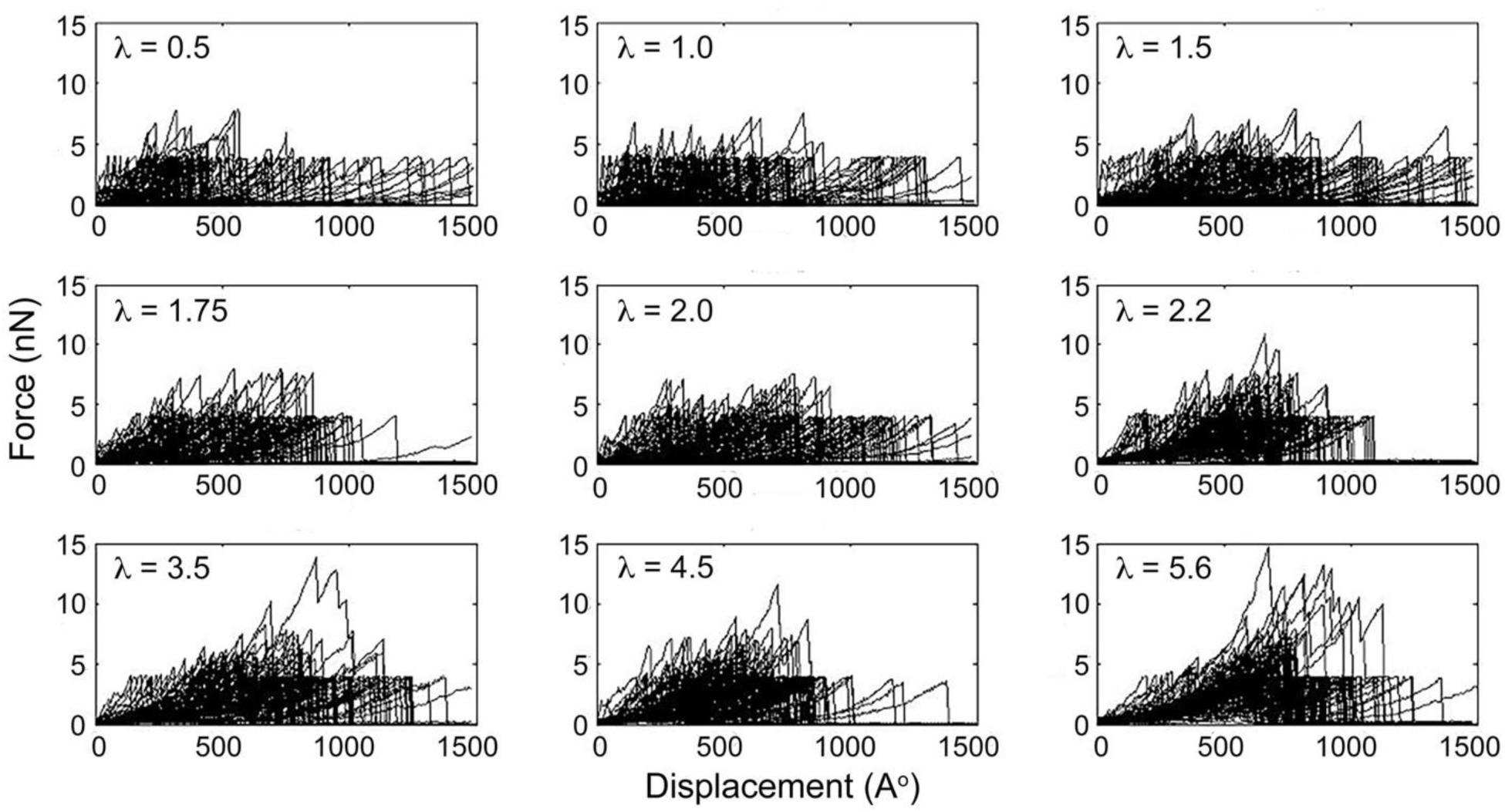
FE curves from MD simulations of pushing single lamin filaments in meshworks of increasing power values (λ = 0.5 – 5.6). Lamin filaments were pushed *in silico* with a constant velocity (0.1 mm s^−1^). Meshwork of each λ value was generated randomly, and thereafter independently tested for generating the FE curves 100 individual times. All the FE curves for each λ value are shown in each panel (λ = 0.5, 1.0, 1.5, 1.75, 2.0, 2.2, 3.5, 4.5 and 5.6).

**Supplementary Fig. 18.**
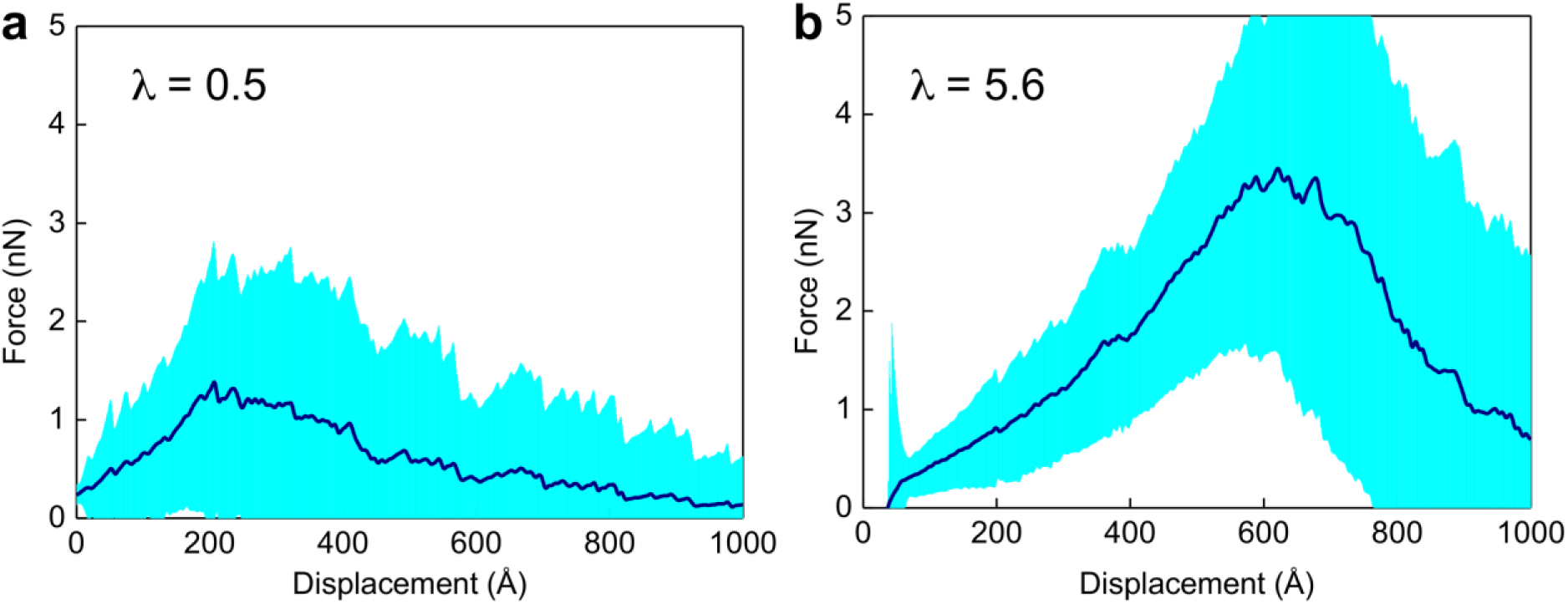
Average force and displacement of lamin filaments depends on meshwork connectivity. (**a, b**) Superimposition of simulated FE curves (turquoise) from meshworks with λ = 0.5 and 5.6, respectively. The average curves (dark blue) clearly show that the filaments can withstand higher forces and deform more in the connected meshwork (λ = 5.6).

**Supplementary Fig. 19.**
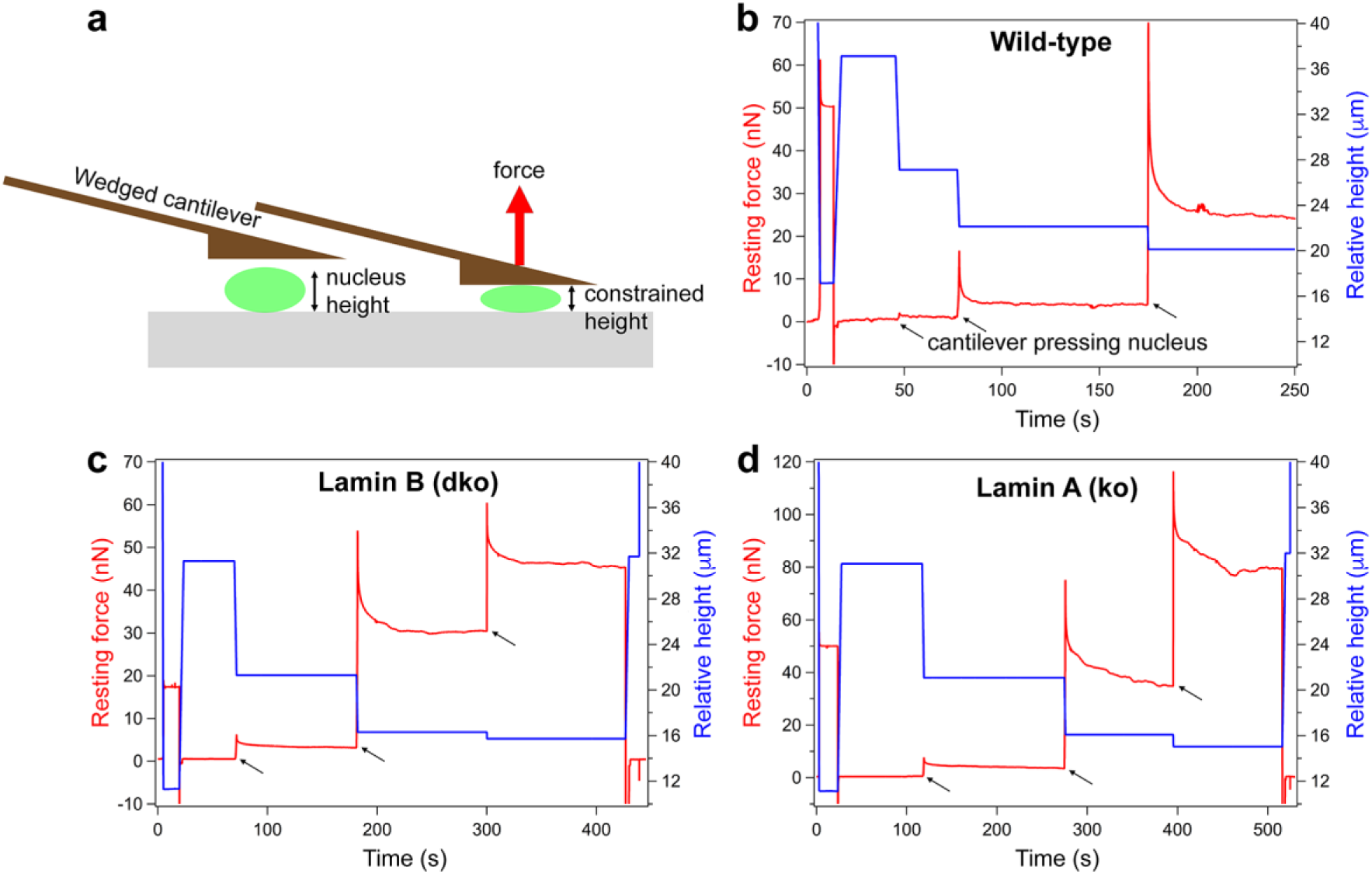
Force resistance of isolated MEF nuclei. (**a**) A single isolated nucleus was subjected to a compression force by confining it between two parallel plates, *i.e.*, between a flat-wedged cantilever and the glass surface. By doing so, the distance between the cantilever and the surface was adjusted such that the nucleus was compressed to 10 μm, 5 μm and 3 μm from its initial height (blue trace) (see **Methods**). (**b – d**) Nuclei were isolated from WT MEFs, lamin B1/B2 double knock-out (dko), lamin A knock-out (ko), constrained to specific heights by the cantilever, and the resting force measured (force reaching steady state after the initial rise, shown by black arrows) (Supplementary Fig. 20). The resting force is the resistance (counterforce) of a nucleus to compression force and may be considered as an indicator of nucleus stiffening. A stiffer nucleus will be able to resist the compressing force more than a softer nucleus. Shown are representative traces from each nucleus type.

**Supplementary Fig. 20.**
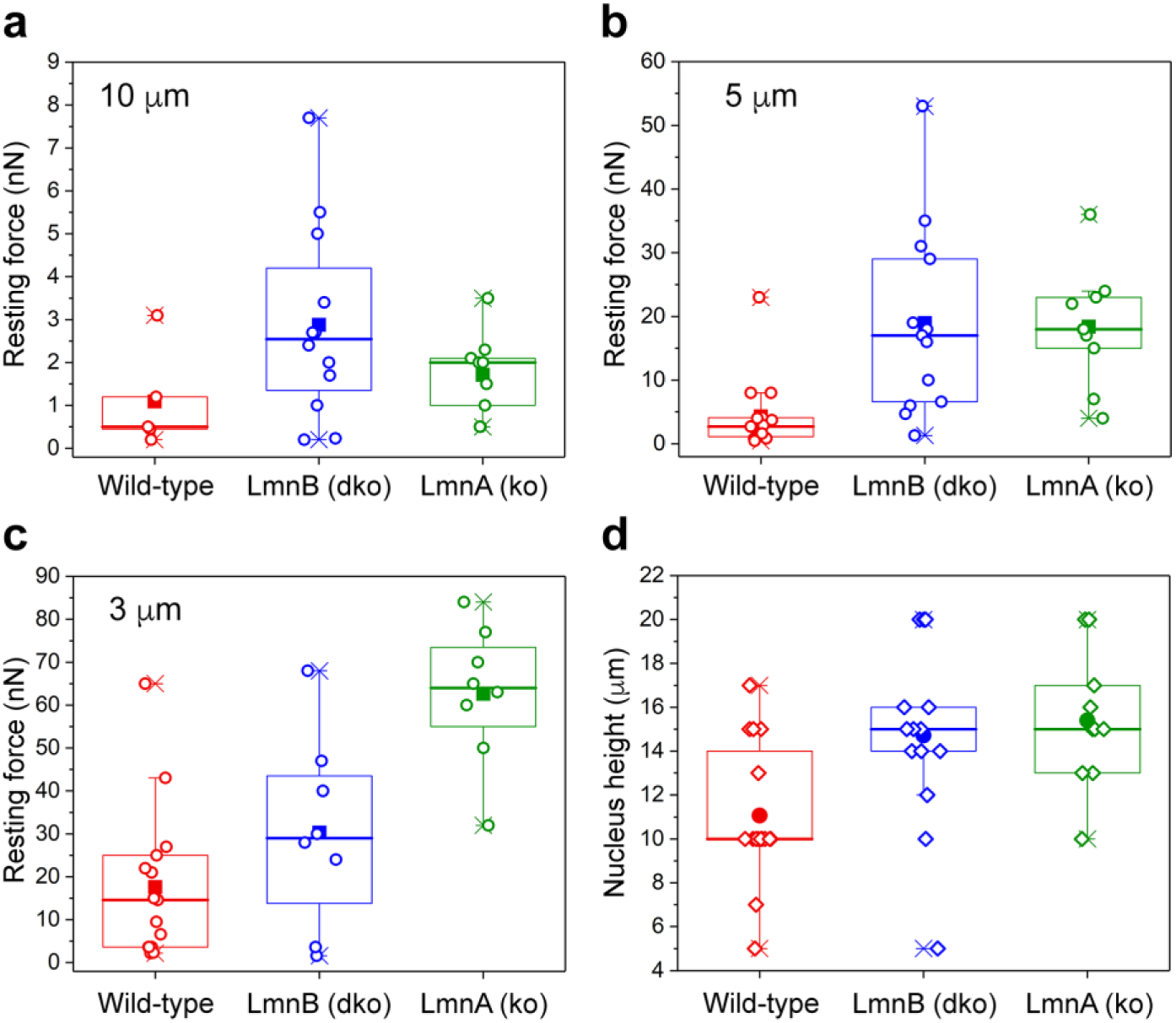
Force resistance of isolated MEF nuclei. Data of resting force experiments of MEF nuclei expressing lamins A/C and B (wild-type), only A-type lamins A/C (Lmn B dko) or only lamin B (Lmn A ko) are plotted. (**a, b**) MEF nuclei expressing only A-type lamins or only lamin B showed higher resting force or increased stiffness compared to wild-type nuclei when compressed to 10 μm and 5 μm heights using AFM cantilevers. (**c**) Upon increasing the compression to 3 μm, lamin B expressing nuclei showed the highest stiffness indicated by an increase in the resistance force by ∼200 % compared to wild-type nucleus. (**d**) Nuclei heights measured before applying compression forces with a flat-wedged cantilever.

**Supplementary Table 1.**
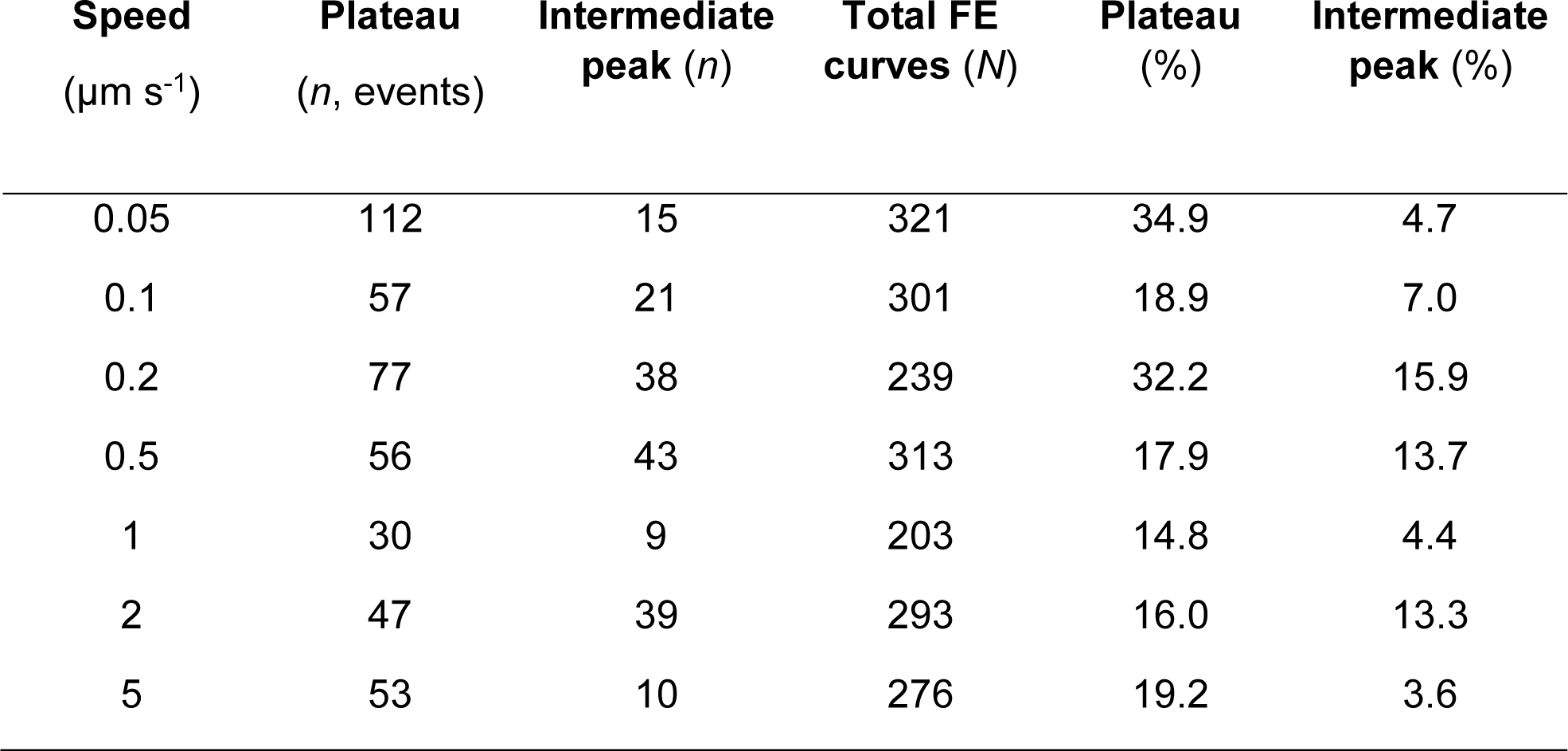
Relative populations of the plateau and the intermediate peak (low force regime) in the FE curves of lamin filaments.

**Supplementary Table 2.** Statistics of the two peaks (step units) from constant velocity force-extension experiments of lamin filaments.

| Peak | Number of events, *n* | Speed ($\mu\text{m s}^{-1}$) |
| --- | --- | --- |
| 1 (all parameters) | 321 | 0.05 |
|  | 301 | 0.1 |
|  | 239 | 0.2 |
|  | 313 | 0.5 |
|  | 203 | 1 |
|  | 293 | 2 |
|  | 276 | 5 |
| 2 (step unit and force) | 187 | 0.05 |
|  | 176 | 0.1 |
|  | 164 | 0.2 |
|  | 215 | 0.5 |
|  | 113 | 1 |
|  | 207 | 2 |
|  | 159 | 5 |
| 2 (stiffness) | 106 | 0.05 |
|  | 104 | 0.1 |
|  | 74 | 0.2 |
|  | 121 | 0.5 |
|  | 43 | 1 |
|  | 144 | 2 |
|  | 100 | 5 |

**Supplementary Table 3.** Relative population of the peaks observed in FE curves in constant velocity experiments.

| **Speed**<br>**($\mu\text{m s}^{-1}$)** | **1<sup>st</sup> peak**<br>**% ± error**<br>**(*n* = )** | **2<sup>nd</sup> peak** | **3<sup>rd</sup> peak** | **4<sup>th</sup> peak** | **5<sup>th</sup> peak** |
| --- | --- | --- | --- | --- | --- |
| 0.05 | 1<br>(321) | $0.6 \pm 0.03$<br>(187) | $0.03 \pm 0.01$<br>(11) | - | - |
| 0.1 | 1<br>(301) | $0.6 \pm 0.03$<br>(176) | $0.04 \pm 0.01$<br>(12) | - | - |
| 0.2 | 1<br>(239) | $0.7 \pm 0.03$<br>(164) | $0.1 \pm 0.02$<br>(24) | $0.02 \pm 0.01$<br>(4) | $0.004 \pm$<br>$0.004$ (1) |
| 0.5 | 1<br>(313) | $0.7 \pm 0.03$<br>(215) | $0.08 \pm 0.02$<br>(24) | $0.01 \pm 0.006$<br>(3) | - |
| 1 | 1<br>(203) | $0.6 \pm 0.03$<br>(113) | $0.02 \pm 0.01$<br>(4) | $0.005 \pm$<br>$0.005$ (1) | - |
| 2 | 1<br>(293) | $0.7 \pm 0.03$<br>(207) | $0.08 \pm 0.02$<br>(23) | $0.02 \pm 0.008$<br>(5) | $0.003 \pm$<br>$0.003$ (1) |
| 5 | 1<br>(276) | $0.6 \pm 0.03$<br>(159) | $0.01 \pm 0.006$<br>(3) | - | - |
| Total | 1<br>(1946) | $0.6 \pm 0.01$<br>(1221) | $0.05 \pm 0.005$<br>(101) | $0.007 \pm$<br>$0.002$ (13) | $0.001 \pm$<br>$0.0007$ (2) |

